# Single cell analysis of senescent epithelia reveals targetable mechanisms promoting fibrosis

**DOI:** 10.1101/2022.03.21.485189

**Authors:** Eoin D O’Sullivan, Katie J Mylonas, Rachel Bell, Cyril Carvalho, David P Baird, Carolynn Cairns, Kevin M Gallagher, Ross Campbell, Marie Docherty, Alexander Laird, Neil C Henderson, Tamir Chandra, Kristina Kirschner, Bryan Conway, Gry H. Dihazi, Michael Zeisberg, Jeremy Hughes, Laura Denby, Hassan Dihazi, David A Ferenbach

## Abstract

Progressive fibrosis and maladaptive organ repair result in significant morbidity and millions of premature deaths annually. Senescent cells accumulate with ageing and after injury and are implicated in organ fibrosis, but the mechanisms by which senescence influences repair are poorly understood. Using two murine models of injury and repair we show that obstructive injury generates senescent epithelia which persist after resolution of the original injury, promote ongoing fibrosis and impede adaptive repair. Depletion of senescent cells with ABT263 reduces fibrosis in reversed ureteric obstruction and after renal ischaemia-reperfusion injury. We validate these findings in humans, showing that senescence and fibrosis persist after relieved renal obstruction. We next characterise senescent epithelia in murine renal injury using single cell RNA-Seq. We extend our classification to human kidney and liver disease and identify conserved pro-fibrotic proteins which we validate in vitro and in human disease. We demonstrate that one such molecule, Protein Disulfide Isomerase Family A Member 3 (PDIA3), is essential for TGF-beta mediated fibroblast activation. Inhibition of PDIA3 in vivo significantly reduces kidney fibrosis during ongoing renal injury and as such represents a new potential therapeutic pathway. Analysis of the signalling pathways of senescent epithelia connects senescence to organ fibrosis, permitting rational design of anti-fibrotic therapies.

## Introduction

Effective repair following injury is essential for maintaining organ function and health. Kidneys are susceptible to injury during illness with acute kidney injury (AKI) complicating up to 25% of hospital admissions (1). Maladaptive repair can lead to scarred kidneys and chronic kidney disease (CKD) (2, 3). Furthermore, CKD itself, a disease affecting almost 700 million people worldwide, increases the risk of AKI and maladaptive repair, which can lead to a vicious cycle of acute injury and disease progression (2, 4). Hence, there is an urgent need to understand the mechanisms of maladaptive repair following injury and develop novel interventions and treatments.

Cellular senescence is a heterogeneous phenotype assumed by cells during development and in response to age, injury or stress. Senescence is an important physiological defence against malignant transformation, and plays an important role in wound healing (5, 6). Senescent cells undergo permanent growth arrest and accumulate in multiple organs with increasing age and disease (7). Senescent cells can adopt a secretory phenotype (known as the Senescence Associated Secretory Phenotype, or SASP). In some contexts, such as wound healing, the SASP can have a beneficial effect, but when chronically present the SASP can alter the tissue microenvironment, exerting a detrimental effect on surrounding cells and ultimately disrupting organ function.

The elderly and those with CKD have the highest burden of senescent cells and are also the most susceptible to maladaptive repair post injury (8). We have previously demonstrated that removal of pre-existing senescent cells prior to kidney injury is protective (9). However, the role of emergent senescent cells after acute injury and during repair remains unknown.

Whilst multiple cell types in the kidney can express markers of permanent cell cycle arrest – it is renal epithelial cell senescence that correlates most strongly to progressive fibrosis and functional loss in human biopsy samples (10, 11). Increased senescent cell epithelial numbers are found in renal biopsies from older patients or those with kidney disease and a greater epithelial senescent cell burden is associated with worse clinical outcomes in several renal conditions including renal transplantation and glomerulonephritis (10, 12–16). Unlike other cell types epithelial cell senescence has been shown to accumulate with advancing age in in murine models of aging and injury (9, 17). The importance of renal epithelial senescence is supported by data which demonstrate that a variety of injuries can induce renal tubular epithelial cells into senescence (18).

For these reasons, whilst not discounting the potential importance of senescence in other cell types within the kidney, we chose to address the role of epithelial senescence in maladaptive repair after resolved injury. We demonstrate that senescent epithelia persist in human kidneys alongside with persistent fibrosis following the resolution of ureteric obstruction. Using experimental models of reversible ureteric obstruction and renal ischaemia reperfusion injury in mice we show that depletion of senescent cells in the aftermath of injury promotes complete repair. In contrast, depletion of senescent cells exacerbated fibrosis during the ‘initiation’ phase of obstructive injury. We characterise senescent epithelial transcriptomes at a single-cell resolution in vivo and explore conserved senescence transcripts across organs and species to identify important profibrotic molecules. One such molecule, Protein Disulfide Isomerase Family A Member 3 (PDIA3), is shown to potentiate TGF-beta mediated fibrosis in vivo. We demonstrate that PDIA3 inhibition during obstructive renal injury is anti-fibrotic in vivo.

## Results

### Senescent epithelia drive fibrosis during post injury renal repair

To explore the role of senescent cells in acute renal injury we first depleted senescent cells using the senolytic drug ABT-263 following unilateral ureteric obstruction (UUO) in mice (Figure 1a). This is a persistent injury model where the insult remains present and tissue damage persists so long as the ureter remains obstructed. Administration of ABT-263 resulted in an increase in renal fibrosis following UUO injury measured by picrosirius red, and a decrease in qPCR markers of senescence (Figure 1b-d). Hypothesising that senescence may thus be important in limiting ongoing injury, we next assessed whether senescent cell depletion following a single discrete injury would have a different effect. We administered ABT-263 to mice following ischemia reperfusion injury commencing at post injury day 3 (Figure 2a) Administration of ABT-263 resulted in reduced kidney fibrosis and improved function by measured Cystatin C clearance in kidneys post IRI (Figure 2b-d). These results suggested senescence may be a desirable phenomenon during ongoing tissue injury but harmful during post injury repair.

**Figure 1.**
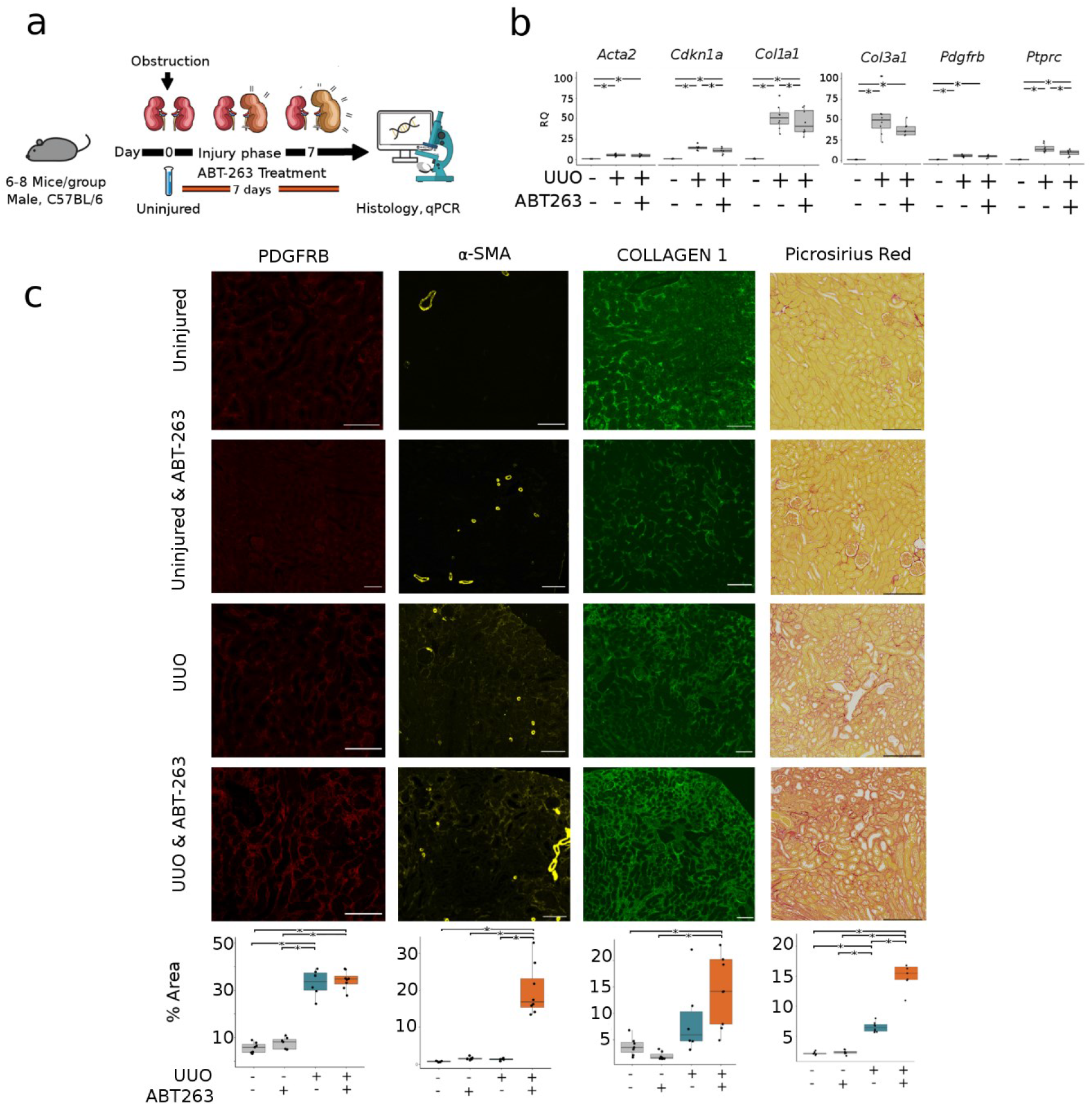
Senescent cell depletion during UUO results in increased renal fibrosis. a) Schematic of murine senescent cell depletion during UUO experiment, with samples taken at days 0, and 7. Note senolytic ABT-263 administration daily from time of injury. b) qPCR analysis of renal tissue post UUO showing reduction in markers of senescence (*Cdkn1a*), no change in myofibroblast markers *Acta2*, *Ptprc*, *Pdgfrb* or *Col3a1*\* denotes P val <0.05 by ANOVA and Tukey test. RQ = relative quantification (fold change) c) Representative images and quantification immunofluorescence and of picrosirius red staining of kidneys. Quantification of total staining per renal cortex. * denotes P val <0.05. Pdgfrb: uninjured & vehicle treated 5.69% (s.d. 2.23), uninjured & ABT263 treated 7.53% (s.d. 2.48). UUO & Vehicle treated 33% (s.d. 5.67) vs UUO & ABT263 treated 34.3% (s.d. 3.77). ANOVA, *P*=0.921 95% CI: -4.2-6.75 a-SMA: uninjured & vehicle treated 0.72% (s.d. 0.16), uninjured & ABT263 treated 1.56% (s.d. 0.39). UUO & Vehicle treated 1.31% (s.d. 0.335) vs UUO & ABT263 treated 19.9% (s.d. 6.96). ANOVA, *P*=<0.000 95% CI: 12.33-24.8 Collagen 1: uninjured & vehicle treated 3.75% (s.d. 1.73), uninjured & ABT263 treated 1.95% (s.d. 0.74). UUO & Vehicle treated 8.85% (s.d. 7.08) vs UUO & ABT263 treated 14.5% (s.d. 6.58). ANOVA, *P*=0.921 95% CI: -4.2-6.75 Picrosirius Red: uninjured & vehicle treated 2.1% (s.d. 0.26), uninjured & ABT263 treated 2.33% (s.d. 0.34). UUO & Vehicle treated 6.5% (s.d. 0.81) vs UUO & ABT263 treated 15.6% (s.d. 3.08). ANOVA, *P*=<0.000 95% CI: 6.8-11.3 For boxplots, the centre line represents the mean, the box limits the first and third quartiles, the whiskers +- 1.5 * IQR and the points all the data. Scale bar = 100 microns.

**Figure 2.**
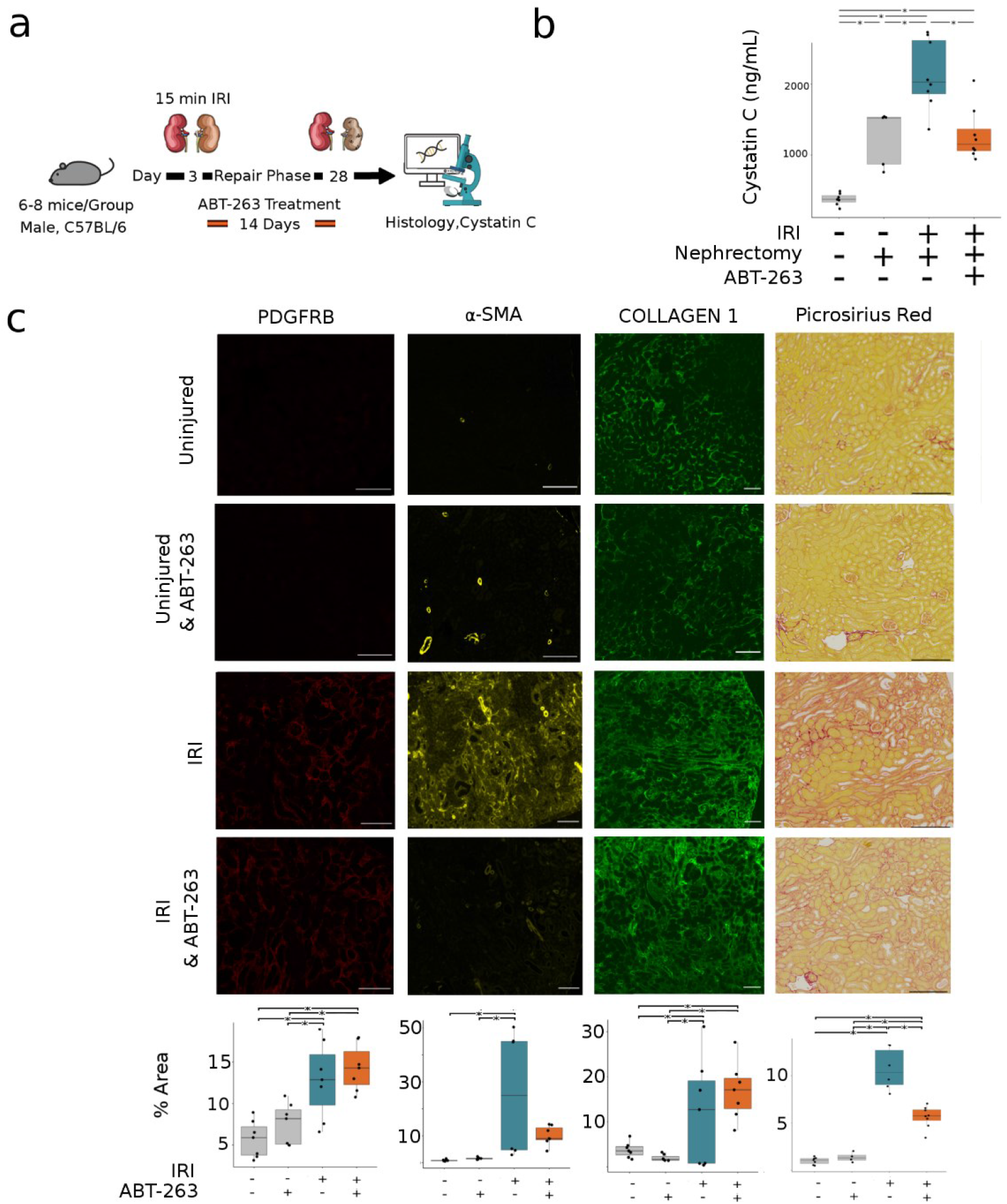
Senescent cell depletion following IRI results in decreased renal fibrosis. a) Schematic of murine senescent cell depletion following IRI experiment, with samples taken at days 0, and 28. Note senolytic ABT-263 administration for 2 weeks following injury. b) Cystatin C measurement in ng/mL – Baseline group, mean Cystatin C 377 (s.d. 84), post nephrectomy group, mean 1244 (s.d. 395). IRI & Vehicle treatment group, mean 2136 (s.d. 491) vs IRI & ABT263 treatment mean 1286 (s.d.372), two-sided t-test, P =0.0006, difference 850, CI: 315-1384. c) Representative images and quantification immunofluorescence and of picrosirius red staining of kidneys. Quantification of total staining per renal cortex. * denotes P val <0.05. Pdgfrb: uninjured 5.69% (s.d. 2.23), uninjured & ABT263 treated 7.53% (s.d. 2.48). IRI & Vehicle treated 12.8% (s.d. 4.66) vs IRI & ABT263 treated 14.3% (s.d. 2.82). ANOVA, *P*=0.921 95% CI: --4.24-6.7 a-SMA: uninjured 0.84% (s.d. 0.33). uninjured & ABT263 treated 1.56% (s.d. 0.39). IRI & Vehicle treated 25.5% (s.d. 23.4) vs IRI & ABT263 treated 10.1% (s.d. 3.56). ANOVA, *P*=0.108 95% CI: -33.74-2.92 Collagen 1: uninjured & vehicle treated 3.75% (s.d. 1.73), uninjured & ABT263 treated 1.95% (s.d. 0.74). UUO & Vehicle treated 12% (s.d. 12) vs UUO & ABT263 treated 16.9% (s.d. 6.39). ANOVA, *P*=0.921 95% CI: -4.2-6.75 Picrosirius Red: control & vehicle treated 0.98% (s.d. 0.37), control & ABT263 treated 1.35% (s.d. 0.51). IRI & Vehicle treated 10.6% (s.d. 2.2) vs IRI & ABT263 treated 5.6% (s.d. 1.14), difference 5.06%, (C.I 3.1-7.01) P=<0.005.For boxplots, the centre line represents the mean, the box limits the first and third quartiles, the whiskers +- 1.5 * IQR and the points all the data. Scale bar = 100 microns. d)

To further explore whether senescent epithelial cells promote maladaptive repair and fibrosis following removal of an injurious stimulus, we used murine reversible unilateral ureteric obstruction (R-UUO), where a single ureter was surgically obstructed before reversal 7 days later. This experimental model is analogous to reversed obstructive uropathy in humans. To establish the role of senescence in maladaptive repair, ABT-263 was administered during the repair phase after injury and compared to a vehicle treated group. Bulk RNA-Seq was performed on uninjured kidneys and 35 days after reversal (“day 42” in figure 3a) to identify the transcriptomic pathways which might prevent full repair and the effect of senolytic treatment.

**Figure 3.**
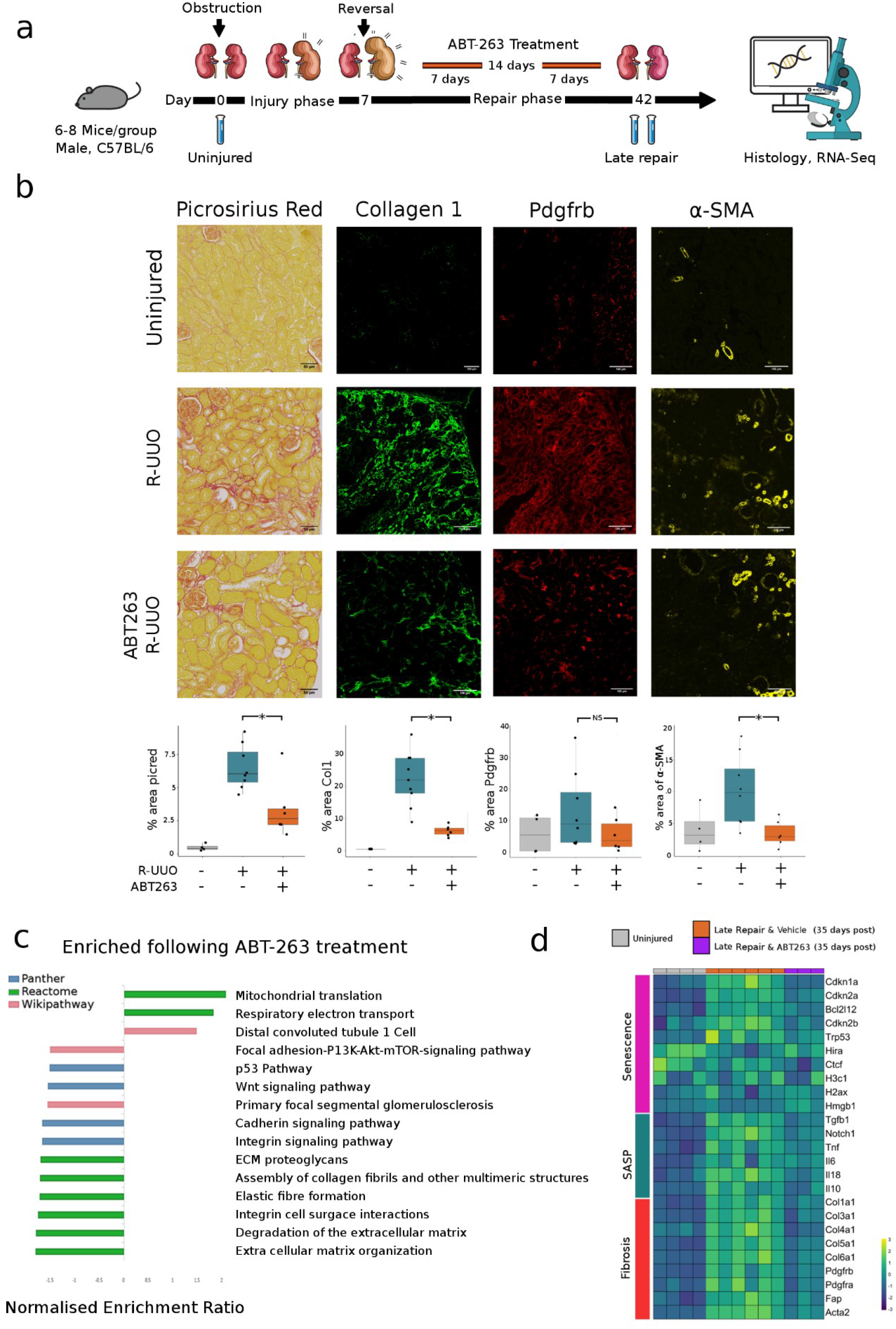
ABT-263 reduces fibrosis and alters the renal transcriptome by depleting senescent cells post R-UUO. a) Schematic of murine R-UUO experiment, with samples taken at days 0, 14 (7 days following reversal) and 42 (35 days following reversal). Note senolytic ABT-263 administration for 14 days during the repair phase. b) Representative images and quantification of picrosirius red staining of kidneys and immunofluorescence of key proteins. Quantification of total staining per renal cortex. * denotes P val <0.05. Picrosirius Red: vehicle treated 6.5% (s.d. 1.6) vs ABT-263 treated 3.3% (s.d. 2.2), two-sided t-test, *P* =0.01, CI: 0.7-5.6. Collagen 1: vehicle treated 5.7% (s.d. 1.6) vs ABT-263 treated 21.5% (s.d. 8.3), two-sided t-test, *P* =0.0003, CI: 9.2-22.3. PDGFRB: vehicle treated 13.1% (s.d. 12.2) vs ABT-263 treated 5.5% (s.d. 5.5), two-sided t-test, *P*=0.1, CI: -3.3-18. α-SMA: vehicle treated 10.2% (s.d. 5.5) vs ABT-263 treated 3.4% (s.d. 2) two-sided t-test, *P* =0.01, 95% CI: 2-11.5. c) Gene Set Enrichment Analysis of DEG between ABT-263 treated animals and vehicle treated controls at 35 days post reversal. All FDR values for all pathways shown are <0.05. GSEA performed on WebGestalt. d) Heatmap showing senescence and fibrosis associated genes are persistent during repair, which return towards baseline following administration of ABT-263. The colour scheme is based on z-score distribution. All genes during late repair are significant DEG (Q val <0.1) as compared to uninjured group. ABT-263 treated genes compared to vehicle treated were all were significant DEG (Q val <0.1) except *H3c1, H2ax, Hmgb1, Il6* and *Il10*.

Mice treated with ABT-263 had significantly less cortical fibrosis as measured by picrosirius red staining and Collagen I immunofluorescence, and reduction of fibroblast activation by α-SMA immunofluorescence (Figure 3b). ABT-263 resulted in a decrease in senescent cells as measured by SA-βgal staining and P21CIP1+ tubules (Supplemental Figure 1a-c). There were marked transcriptional changes during repair and following ABT-263 treatment (Figure 3c). Gene set enrichment analysis revealed pathways associated with persistent inflammation and fibrosis including interleukin signalling and extracellular matrix reorganisation remained persistently elevated after reversal of obstruction (Figure 3c). At day 35 post reversal, mice treated with ABT-263 had a significant down-regulation of pathways associated with senescence, the SASP fibrosis and including P53 pathway, extracellular matrix organisation and elastic fibre formation when compared to vehicle treated mice (Figure 3d, supplemental table 1).

Taken together, these data indicated that senescent epithelia persisting in the aftermath of injury promote maladaptive repair, and the depletion of senescent cells by ABT-263 in this setting reduces subsequent renal fibrosis and mesenchymal activation following de-obstruction.

### Senescent epithelial cells are associated with post injury fibrosis in human kidneys

Having observed persistence of senescence in murine epithelia following injury we next examined whether senescent cells induced during a similar obstructive kidney injury also persisted in humans despite resolution of injury. To do this we utilised the non-tumorous portion of tumour nephrectomy specimens from three cohorts of patients; 1) kidneys with no evidence of obstruction, 2) persistently obstructed kidneys and 3) previously obstructed kidneys where obstruction had been relieved by prior placement of a ureteric stent (Figure 4a). Patients who had persistent obstruction had increased levels of fibrosis and expression of the cyclin dependent kinase inhibitor P21CIP1 (typically upregulated in senescence) (Figure 4b,4c). Notably, P21CIP1 expression and fibrosis levels did not return to baseline post obstruction, suggesting persistent senescence may be present alongside maladaptive repair even after relief of obstruction (Figure 4b, 4c).

**Figure 4.**
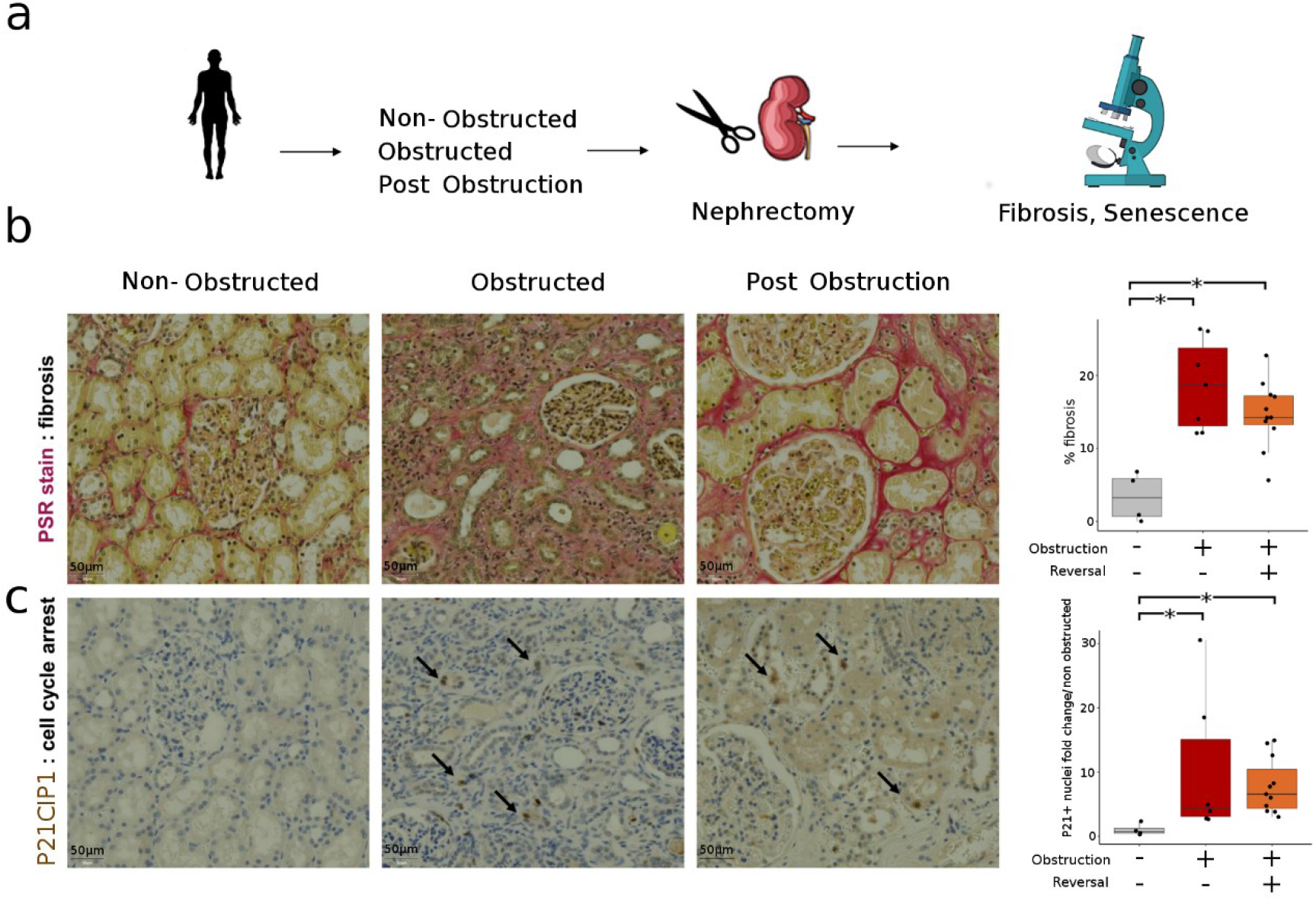
Senescent cells persist following renal injury with subsequent maladaptive repair. a) Schematic of human kidney sample retrieval (n=22) b) Human kidney fibrosis. Red staining indicates picrosirius red staining of collagen networks. * denotes significance at P val<0.05. Uninjured 3.3% (s.d. 3.3) vs obstruction 18.6% (s.d. 6.1) vs post obstruction 14% (s.d. 4.5), ANOVA, uninjured vs obstructed, *P* =0.002 CI 7.4-23.3, uninjured vs post obstruction *P* =0.002 CI 3.9-18.7, obstruction vs post obstruction *P* =0.6 CI -10.14-2.11. For all boxplots, the centre line represents the mean, the box limits the first and third quartiles, the whiskers are +- 1.5 * IQR. c) Human kidney senescence. P21CIP1 staining demonstrating senescent epithelial cell. Arrows point to P21CIP1 positive nuclei. Y axis shows fold change of senescent tubules per kidney relative to the mean number of senescent tubules in the uninjured control group. Uninjured 0.9 (s.d. 0.9) vs obstruction 10.5 (s.d. 11.4) vs post obstruction 7.8 (s.d. 4.3), Kruskal-Wallis rank sum test, uninjured vs obstructed, *P* =0.009 CI 1.5-29.6, uninjured vs post obstruction *P* =0.001 CI 2.7-12.5, obstruction vs post obstruction *P* =0.2 CI -5.5-14.5.

### Single-cell RNA-Seq allows analysis of senescent cells in vivo

To better understand how senescent cell persistence inhibits repair and promotes fibrosis we examined subsequent studies using single-cell RNA-Seq. Single-cell libraries were prepared using whole kidney digests of three pooled mice per library at multiple time points: day 2 post UUO, day 7 post-UUO and 14 days post-R-UUO (Figure 5a). This dataset was recently published analysing the myeloid compartment (19). All cells can be explored via our online atlas http://www.ruuo-kidney-gene-atlas.com. As we have previously described the varied cell types during R-UUO in detail, and as senescent renal epithelia are implicated in the induction of fibrosis after injury as discussed, we focused this analysis on the epithelial compartment.

**Figure 5.**
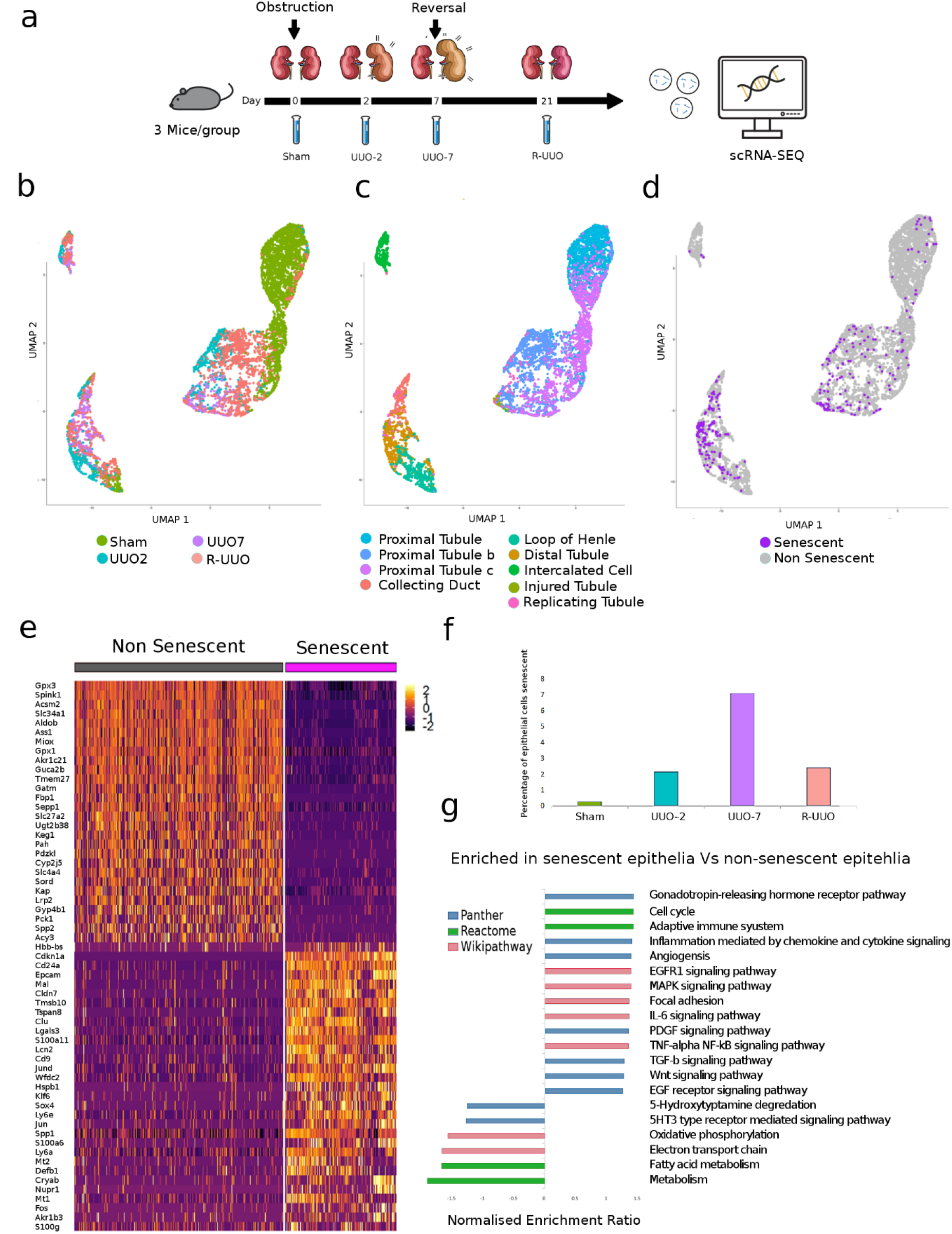
Senescent cells exhibit a proinflammatory, profibrotic transcriptome. a) Schematic of single cell RNA-Seq R-UUO experimental workflow UMAP plots of 7392 epithelial cells coloured by b) injury timepoint; c) unbiased cell cluster and d) by senescent status (320 cells). e) Heatmap of top DE genes between non/senescent epithelial cells, calculated using Wilcoxon signed-rank test. The colour scheme is based on z-score distribution. f) Percentage of epithelial cells classified as senescent at each injury timepoint. The proportion of epithelial cells which are senescent increases over time following injury and decreases during repair but does not return to baseline. g) Gene Set Enrichment Analysis between senescent and non-senescent epithelial cells. Selected pathways demonstrate consistency with known senescent signalling pathways. All FDR values for pathways shown are <0.05. GSEA scores generated with WebGestalt as described in methods.

Epithelial cells were first identified following unsupervised clustering and classification before epithelial sub-types (e.g, proximal tubule, loop of Henle etc) were classified based on characteristic marker genes (Figure 5b, 4c, Supplemental Figure 1d,1e,2a,2b). An “injured tubule” cluster was identified and uniquely characterised by elevated injury marker *Havcr1* (KIM-1), which was transcriptomically distinct from other clusters (Figure 5c). Next, to classify a subgroup as senescent, we over-partitioned the epithelia into multiple clusters. A single cluster was identified expressing high *Cdkn1a* and an absence of cells in S phase - and was thus classified as senescent. These cells demonstrated upregulation of typical senescent pathways (SASP genes, TGFb and Notch signalling) (Supplemental Figure 3, 4).

In total, 322 of the 7392 (4.2%) epithelial cells were classified as senescent (Figure 5d). Senescent cells had a markedly different transcriptome, accumulated over time following injury and decreased following de-obstruction but did not return to baseline, consistent with our earlier observations in post obstruction human kidneys (Figure 5e-g, supplemental table 2). Senescent cells were most frequently found amongst distal tubular cells, consistent with previous observations in human renal senescence (20).

### Senescent cells exhibit a pro-fibrotic, pro-inflammatory transcriptome

When compared to non-senescent epithelial cells, the first notable pattern amongst the differentially expressed genes in senescent epithelial cells was the loss of normal tubular function (Figure 5e). This apparent “dedifferentiation” remained when comparing senescent cells only to the distal tubular cell cluster, indicating that this phenomenon was not due to any senescent cell bias towards distal tubule or transcriptomic differences along the tubule. Senescent cells expressed typical SASP components as well as being enriched for transcripts of *Notch* and *Tgf* signalling pathways (Supplemental Figure 4) (21, 22).

Gene set enrichment analysis showed enrichment for multiple relevant ontologies (Figure 5g) including: IL6 signalling, an important component of the SASP,(23, 24) TGFβ signalling, an inducer of senescence and renal fibrosis; WNT signalling, crucial in cell fate decisions by regulating p53,(25) and PDGF signalling, important in myofibroblast differentiation and underpinning senescence in healthy wound healing (26–29).

To validate our transcriptomic signature in other dataset sets, we used a machine learning classifier which we trained on our signature and then used to classify cells in three public datasets(30). Senescent cells were classified in publicly available datasets - two ischemia reperfusion injury models and “tabula muris senis”, which includes aged mice. We successfully identified senescent cells in all these data and found similar enriched pathways in all cases (supplemental figure 5). To corroborate our findings we compared the differentially expressed genes (DEGs) in our senescent epithelial cells with the proteome of senescent human cortical renal epithelial cells in the “SASP Atlas”(31). We found 363/860 (42%) of the proteins found in the SASP Atlas appeared as DEGs in our senescent epithelial cell DEG list. These data support the hypothesis that senescent epithelial cells produce a profibrotic, inflammatory SASP. Next, we used two analytical approaches to identify candidate profibrotic molecules for further validation.

### Single-cell RNA-Seq reveals conserved senescent gene expression between organs and species

First, considering senescence is a highly conserved biological cell phenotype we reasoned that DEGs in senescent epithelial cells that are observed across species and organs may be important pathogenic molecules representing good candidates for further interrogation as therapeutic targets. We therefore reanalysed single cell libraries from a human kidney transplant from a 70 year old donor where the patient developed chronic allograft nephropathy, as we anticipated this would include chronic senescent cells due to both donor age and ongoing disease (32). Of the 4487 epithelial cells in the single-cell RNA-Seq dataset, 114 cells (2.5%) were classified as senescent (Figure 6a). Next, we explored recently published single cell RNA-Seq from cirrhotic human livers and matched control tissue and we used the same approach to senescent cell identification described above (33). Of the 3689 epithelial cells in this dataset, 134 senescent epithelial cells were identified (3.5% total), 130 of which were derived from the cirrhotic library and 4 cells from the healthy liver (Figure 6b). Importantly, several senescent epithelial DEGs were found to be shared across both human datasets and our murine dataset (Figure 6c).

**Figure 6.**
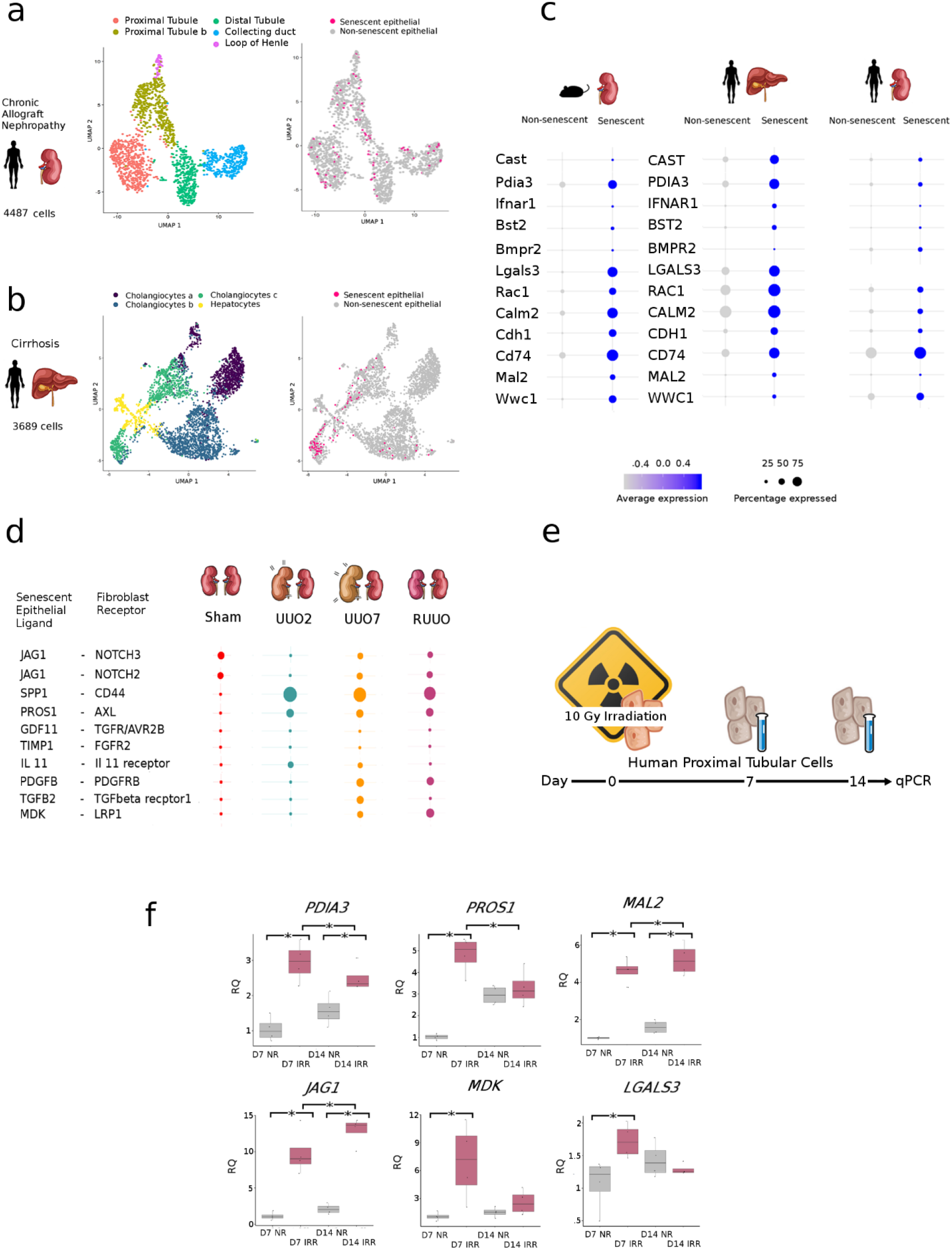
Conserved senescent transcriptome across species and organ allows identification of potential therapeutic targets. a) UMAP of 3869 epithelial cells from human cirrhotic livers, coloured by classification from original paper, and senescent status. b) UMAP of 1692 epithelial cells from a patient with chronic allograft nephropathy coloured by cell type classification and senescent status c) Expression of DEG in senescent epithelial cells which were consistent across species and organs. Size of dot represents the percentage of cells in each cluster expressing the gene, colour indicates average gene expression per cell. d) Top Senescent ligand-mesenchymal receptor pairs across the time-course of the reversible ureteric obstruction model. Size of dot is proportional to mean expression values for all the interacting partners. All interactions are significant with P-val< 0.05. P-value refers to the enrichment of the interacting ligand-receptor pair in each of the interacting pairs of cell types. e) Experimental workflow for inducing senescence in irradiated human proximal tubular cells. f) Irradiation induced senescence in human proximal tubular cells increases key ligand transcription. IRR= irradiated, NR=no irradiation. ANOVA test used for significance testing, * denotes P val <0.05. For boxplots, the centre line represents the mean, the box limits the first and third quartiles, the whiskers +- 1.5 * IQR and the points all the data.

### Single-cell RNA-Seq identifies candidate ligand/receptor interactions between senescent epithelia and fibroblasts during repair

Secondly, based on our earlier data which demonstrated persistent mesenchymal activation during maladaptive repair with enriched pathways in senescent epithelia including TGF-β and PDGF-β (Figure 5g), we analysed senescent epithelia-myofibroblast crosstalk using ligand-receptor pair analysis (34). We found that senescent epithelial cell ligand-fibroblast receptor interactions varied over the course of injury and repair (Fig 6d, myofibroblast classification described in Supplemental Figure 6).

Combining our data from a) conserved DEGs in multi-organ and species senescence and b) ligand-receptor analysis we selected seven candidate molecules for further analysis (Supplemental Table 3).

### Candidate profibrotic mediators are upregulated by senescent human proximal tubular epithelial cells in vitro and in chronic kidney disease in vivo

To further confirm that these molecules were produced in excess by senescent epithelial cells as suggested by the in silico data, we induced senescence in human proximal tubular epithelial cells (hPTEC) using irradiation, an established model of senescence induction (Figure 6e). At day 7 following irradiation, compared to non-irradiated, time matched controls, hPTECs showed increased levels of all candidate molecules as measured by qPCR, with *PDIA3*, *MAL2* and *JAG1* transcripts remaining elevated above non-irradiated controls at day 14 (Figure 6f).

Staining of human protein atlas samples (age range 26-70) demonstrated that our candidate molecules were found in human tubular epithelium and upregulated in human CKD (Supplemental Figure 7a), with Protein Disulfide-Isomerase A3 (PDIA3) showing the strongest correlation (FC >2, p=4.4e-19 vs healthy control). PDIA3 is a thiol oxidoreductase known to be induced by endoplasmic reticulum (ER) stress and capable of mediating cellular responses to oxidative stress. The importance of PDIA3 was not restricted to senescent cells but also enriched in other model of stress and injury, with upregulation also found in other tubular cells after murine UUO and in tubular cells of diabetic patients as compared to healthy controls as well as a proteomic atlas of senescent cells supernatant (Supplemental Figure 7b, 7c).

### PDIA3 is essential for fibroblast proliferation and SMAD2 phosphorylation in response to TGFβ1 in vitro

PDIA3 was selected for further study on the basis of its upregulation in the proteomic SASP atlas (31), in human CKD, our in vitro senescence model and in human and murine senescent cells in vivo. In the kidney, fibroblasts are the key cell responsible for matrix deposition and fibrosis and were therefore identified as the most important downstream target of epithelial PDIA3 during maladaptive repair. However, treatment of fibroblasts with recombinant PDIA3 alone did not result in activation (Supplemental Figure 7g). To test whether PDIA3 augmented the fibroblast response to known activating pathways, we used siRNA (siPDIA3) and a *PDIA3* overexpression plasmid (pPDIA3) to knockdown or over-express PDIA3 in human TK188 fibroblasts prior to activating them with TGF-β (Figure 7a, 7b Supplemental Figure 7d). TGF-β increased fibroblast viability, and this effect was augmented by pPDIA3. In combination, TGF-β and pPDIA3 resulted in increased numbers of viable fibroblasts in this assay and on further replication using Trypan blue to assess cell counts (Figure 7b, 7c). There was a loss of fibroblast response to TGF-β following the silencing of *PDIA3* by siPDIA3 (Figure 7b) and after administration of the pharmacological PDIA3 antagonist Loc14 (Figure 7d).

**Figure 7.**
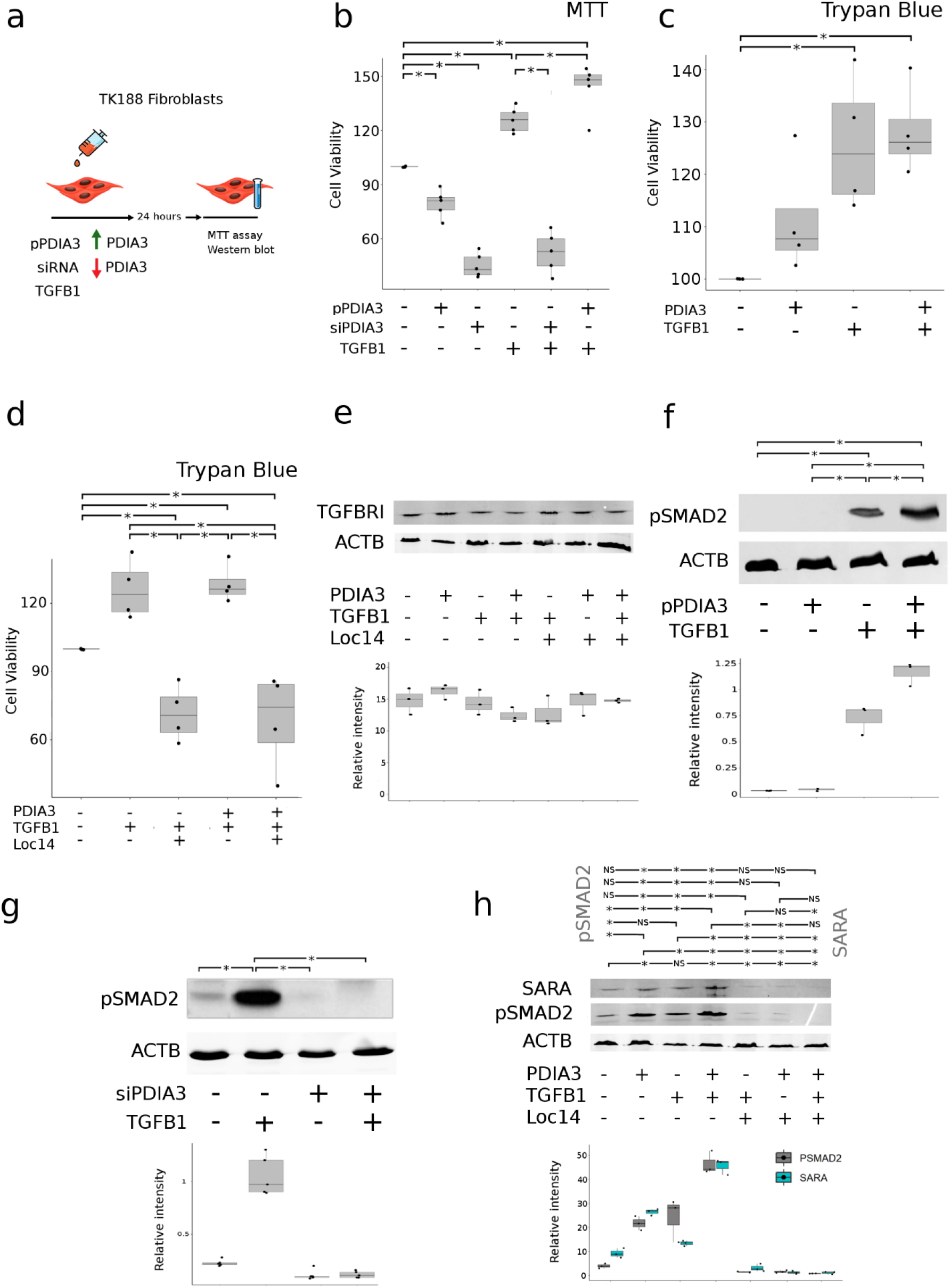
PDIA3 is essential for and enhances TGFB1 driven fibroblast proliferation in vitro. a) Experiment schematic showing modulation of PDIA3 in human fibroblasts qPCR of human kidney fibroblasts treated with PDIA3. b) MTT assay readout normalised to control group demonstrating that dual treatment with pPDIA3 and TGFB1 results in additional proliferation compared to single TGFB1. All comparisons were significant with P adj< 0.05 using one way ANOVA with Bonferroni correction with the exception of siRNA vs siRNA & TGFB1 where P adj = 0.8. Importantly, pPDIA3 & TGFB1 vs TGFB1 showed a difference of 17.8, CI 0.5-35.1, P adj = 0.039. * denotes significance at P val<0.05 c) Trypan blue assay readout normalised to control group demonstrating treatment with PDIA3 and/or TGFB1 results in additional proliferation compared control. Control group mean 100% (s.d. 0) vs TGFB1 mean 125.9% (s.d.12.9), difference 25.9 (C.I -2.3 – 54.24) *P*= 0.01, control vs PDIA3 mean 111.3% (s.d. 11), difference 28.2 (C.I. 8.3-45.9),P=0.005.TGFB1 & PDIA3 mean 128.7% (s.d. 8.5). All comparisons were significant with P adj< 0.05 using one way ANOVA with Bonferroni correction. * denotes significance at P val<0.05 d) Trypan blue assay readout normalised to control group demonstrating treatment with Loc14 reduces cell viability. Control group mean 100% (s.d. 0), control vs TGFB1 & PDIA3 128.27% (s.d. 8.5), difference 28.2 (C.I. -0.01 – 56.5), *P*=0.05. Control vs TGFB1 & Loc14 mean 71.3 (s.d.12.3), difference 28, (C.I 0.36 – 56) *P*=0.04. Control vs TGFB1 & PDIA3 & Loc14 mean 68.7% (s.d. 21.1), difference 31.2 (C.I. 2.9-59.5) *P*=0.02. TGFB1 vs TGFB1 & Loc14 difference 54 (C.I. 26.3-82.9) P=0.0002, TGFB1 vs TGFB1 & PDIA3 & LOC14, difference 57.2 (C.I. 28-85.5) *P*=<0.001. TGFB1 & Loc14 vs TGFB1 & PDIA3, difference 56.9 (C.I. 28.6 – 85), *P*=<0.001. TGFB1 & PDIA3 vs TGFB1 & PDIA3 & Loc14, difference 29.5 (C.I. 31.2-87.8), *P*=<0.001. * denotes P val <0.05 by ANOVA with Bonferroni correction. * denotes significance at P val<0.05. e) Coadministration of PDIA3, TGFB1 and Loc14 to human renal fibroblasts does not alter TGFBRI synthesis. Comparisons tested using one way ANOVA with Bonferroni correction. No comparisons were statistically significant at P adj val<0.05. f) Coadministration of pPDIA3 to TGFB1 treated fibroblasts increases pSMAD2 synthesis. Comparisons tested using one way ANOVA with Bonferroni correction. Significance set at P adj< 0.05 g) Coadministration of siRNA to TGFB1 treated fibroblasts reduces pSMAD2 synthesis. Comparisons tested using one way ANOVA with Bonferroni correction. Significance set at P adj< 0.05 h) Coadministration of PDIA3, TGFB1 to human renal fibroblasts increases SARA and pSMAD2 synthesis. Dual treatment with PDIA3 and TGFB1 results in a greater increase compared to either alone. This effect was abolished by Loc14 administration. Comparisons tested using one way ANOVA with Bonferroni correction. * denotes significance at P adj val<0.05

Next, to test whether this enhanced effect of TGF-β was due to PDIA3 converting latent TGF-β to its active form, an active TGF-β immunoassay was used to test supernatant from renal epithelial cells incubated with latent TGF-b with and without PDIA3 and Loc14, a PDIA3 inhibitor. PDIA3 did not increase free active TGF-β (supplemental figure 7e). Similarly, western blotting demonstrated that total levels of TGFBR1 remained unaltered by Loc14 (Figure 7e). To test the impact of PDIA3 on canonical TGF-β signalling downstream of TGF-β receptor ligation, western blot analysis of pSMAD2 was undertaken. In response to silencing of *PDIA3* during TGF-β treatment, there was diminished phosphorylation of SMAD2 protein, which is required for TGF-β signal transduction. Conversely there was increased phosphorylation of SMAD2 when pPDIA3 was given in combination with TGF-β (Figure 7f, 7g). Further western blotting demonstrated that PDIA3 administration significantly increased expression levels of SARA (Smad anchor for receptor activation) - a cofactor essential for TGF-beta induced Smad2 activation, with levels of SARA potently inhibited by Loc14 treatment (Figure 7h). Collectively, these data demonstrate that PDIA3 plays an essential role in the canonical activation of fibroblasts by the prototypic profibrotic cytokine TGF-β, via intracellular effects on SARA levels, with inhibition of PDIA3 inhibiting TGF-β induced activation in vitro.

### Targeted inhibition of PDIA3 inhibits fibroblast proliferation and fibrosis deposition in vivo

We tested PDIA3 inhibition in vivo in mice during renal injury and repair. An orally bioavailable inhibitor of PDIA3, Loc14, was delivered by gavage to mice following IRI, UUO and R-UUO.

Mice treated with Loc14 during UUO showed significantly less renal fibrosis (Figure 8a). Consistent with our in vitro findings, these mice showed significantly less pSMAD2 staining and less fibroblast proliferation and activation as measured by PDGFRβ and α-SMA immunofluorescence (Figure 8b). This resulted in less fibrosis on picrosirius red staining and Type I Collagen by immunofluorescence (Figure 8d). No toxicity or alteration in levels of renal fibrosis were seen in uninjured control kidneys treated with Loc14 vs vehicle (supplemental figure 7f) and inhibition did not impact senescent cell number, as measured by *Cdkn1a* or *Cdkn2a* via qPCR or P21CIP1 expression on immunofluorescence (supplemental figure 8c, 8d).

**Figure 8.**
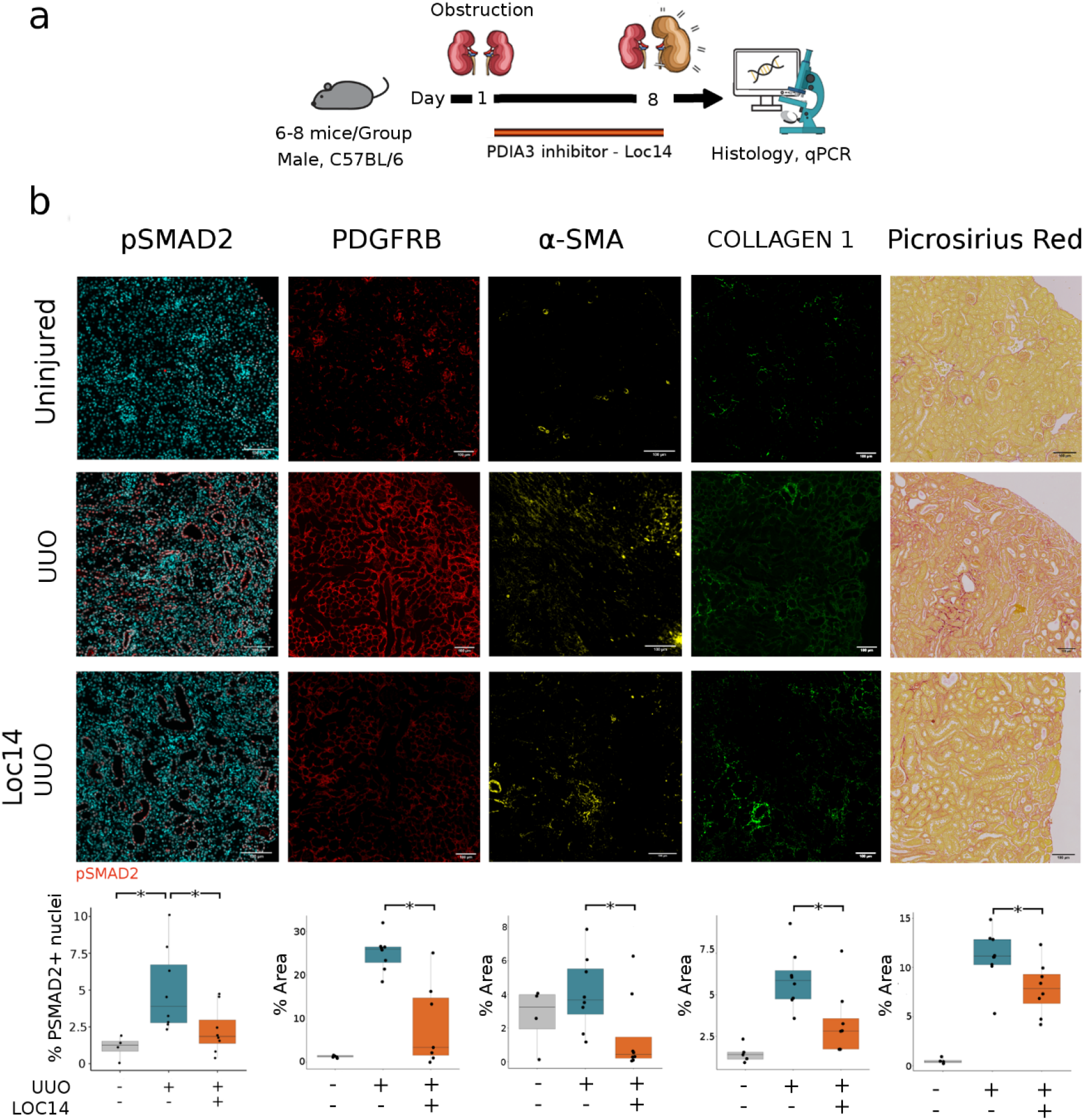
Inhibition of PDIA3 reduces SMAD2 phosphorylation, fibroblast proliferation and fibrosis deposition after UUO *in vivo*. a) Experimental workflow for inhibition of PDIA3 following UUO. d) Representative images and quantification immunofluorescence and of picrosirius red staining of kidneys. Quantification of total staining per renal cortex. * denotes P val <0.05. Loc14 administration results in less pSMAD2 activation in vivo post UUO in mice. Mean% staining in uninjured group 1.89% (s.d. 0.7) vs Vehicle treated post UUO 4.99% (s.d. 2.8), ANOVA, *P adj*=0.02, CI: 0.5-7.2. vehicle treated 4.99% (s.d. 2.8) vs Loc14 treated 2.27% (s.d. 1.6) ANOVA, *P adj* =0.05, 95% CI: -=5.44-0.010. Picrosirius red: 11% (s.d. 2.8) in vehicle treated vs 7.8% (s.d. 2.6) in loc14 treated mice, two-sided t-test, *P* = 0.03, 95% CI: 0.2-6.18. Collagen 1: 3.2% (s.d. 2) in vehicle treated vs 5.9 % (s.d. 1.7) in Loc14 treated, Wilcoxon rank sum test, *P* =0.01, CI 1:4.4. Pdgfrb: 25% (s.d. 4) in vehicle treated vs 8.7% (s.d. 4) in loc14 treated mice, two-sided t-test, *P* = 0.003, 95% CI: 7.2-25. α-SMA: 4% in vehicle treated (s.d. 2.2) vs 1.5% in loc14 treated mice (s.d. 2.3), two-sample Wilcoxon test, *P* = 0.037, CI 0.6-5.2. For boxplots, the centre line represents the mean, the box limits the first and third quartiles, the whiskers +- 1.5 * IQR and the points all the data.

We quantified the nuclei of PDGFRB+ cells and found a statistically significant increase in the injured vehicle treated group as compared to uninjured control animals, with a significant reduction back to baseline levels in Loc14 treated animals. We quantified the nuclei of α-SMA+PDGFRB+ cells and found a statistically significant increase in the numbers of such cells following injury, but no significant change in number following Loc14 treatment. This data is consistent with the primary effect of PDIA3 inhibition being reduction of mesenchymal proliferation rather than reduced myofibroblast activation during acute injury (supplemental figure 7h).

When Loc14 was tested in the reversal phase of R-UUO, it resulted in less Collagen 1 by immunofluorescence, non-significant trends towards decreased markers of activated myofibroblasts and total collagen staining with picrosirius red and unaltered levels of PDGFRB. This is in keeping with a primary role for PDIA3 in augmenting fibrosis deposition rather than preventing its removal (supplemental figure 8a, 8b).

Collectively, these data indicate that PDIA3 facilitates TGF-β mediated fibroblast proliferation in vitro and in vivo via phosphorylation of SMAD2. The effect of its inhibition is context and injury dependant and can reduce fibroblast numbers and collagen production.

## Discussion

The role of persistent senescence in the aftermath of acute injury in facilitating or opposing complete tissue repair remains poorly understood. We show that senescent cells persist in de-obstructed kidneys in experimental and human renal disease and after experimental IRI where they oppose successful renal repair. The effect of experimental depletion of senescent cells is context dependant, and during the repair phase reduces post injury fibrosis. Using single-cell analysis of murine and human kidneys and human fibrotic liver disease we identify factors generated by senescent epithelial cells not previously associated with the senescence-associated secretory phenotype. We characterise one such mediator, PDIA3, as a major contributor to renal fibrosis via its augmentation of TGF-β driven activation of myofibroblasts. We then demonstrate the potential anti-fibrotic efficacy of its targeted inhibition in vivo during ongoing injury.

We provide in silico and in vitro evidence that senescent epithelial cells activate fibroblasts and identify new proteins which may present future therapeutic targets, validating PDIA3 as an example. We hypothesised PDIA3 plays a central role in cell-stress associated fibrosis, which would unify past observations made of the association between PDIA3 and fibrosis in disease settings (35–38). PDIA3 has a number of functions that could be linked to a variety of stressors, including oxidative, nutritional (mTORC1 activator), proteostasis (chaperone function), mitogenic (role in cell division) and oxidative stress. PDIA3 knock out models have been associated with less fibrosis in murine models of lung injury (35, 36) muscle injury (37) and traumatic brain injury (38). PDIA3 is also described in the kidney, where it is upregulated in human CKD and a murine model of CKD (39). Intriguingly, it was demonstrated that ER stress alone is insufficient to induce a secretable form of PDIA3 – where it may have direct ECM stabilisation function, instead requiring a more pro-inflammatory/ SASP like initiation, such as TGF-β. Based on our data, it appears the inhibition of PDIA3 inhibits pSMAD mediated TGF-β intracellular signal transduction. To our knowledge, this is the first inhibition of PDIA3 in kidneys in vivo and provides a connection between the cellular stress response (present in senescent and non-senescent cells) and tissue fibrosis.

There are limitations to a single cell RNA-Seq workflow in this setting. Conventional senescent markers such as p16*^INK4a^* (CDKN2A) and SA-β-gal (GLB1) are poorly sequenced and are often not present in RNA-Seq data of senescent cells. This is compounded by the high dropout intrinsic to single-cell workflows, necessitating a more involved classification process and requiring sensitivity to be weighed against specificity when setting thresholds and classifying cells. We optimised our workflow for specificity, at the natural expense of sensitivity, and as a result have probably under-classified and under-quantified the true number of senescent cells in our data. The cells we have not captured may have a varied transcriptome worth investigating, but the risk of false positives obscuring ligands and DEG outweighed this consideration. LOC14 is a PDI inhibitor and it is possible that LOC14 will react and oxidize both catalytic domains of PDIA1 and PDIA3 (40). It is therefore reassuring that consistent effects were noted with our siRNA studies which are specific to PDIA3. The role of TGFB during injury and repair is complex and it is not surprising that inhibiting a partner molecule results in heterogenous outcomes across injuries. More work is required to understand the optimal clinical scenarios where inhibiting PDIA3 and where depleting senescent cells would be indicated.

In summary, our data shed light on the role of senescent epithelia in renal disease. We demonstrate increased senescence and persistent fibrosis following injury in humans, attenuation of fibrosis following senolysis with ABT-263 in a relevant mouse model and we use single-cell RNA-Seq to describe the in vivo senescent landscape. We identify PDIA3 as a SASP component and an upregulated protein in cells undergoing stress. Furthermore, we link this response with tissue fibrosis and demonstrate that its inhibition reduces fibrosis in a model of kidney disease. Our work demonstrates the ability of single cell transcriptomics to reveal novel senescence and stress associated signalling pathways leading to new, evidence-based anti-fibrotic therapies.

## Methods

All reagents were purchased from Sigma-Aldrich, St Louis, MO, USA unless otherwise stated.

### Human tissue samples

Human kidney samples from three cohorts of patients were analysed. Non-tumour kidney was analysed following nephrectomy for renal cell carcinoma in patients with no obstruction (n=4), obstruction (n=7), and following de-obstruction (n=11). Samples were obtained under local ethical approval, project number SR1405 from patients in the southeast Scotland.

### Mice

Mice were purchased from Charles River laboratories (Wilmington, Massachusetts, United States), Envigo (Envigo, Indianapolis, USA) or bred in house before use. All mice housed in a pathogen-free environment at the University of Edinburgh Animal Facility and were fed standard rodent chow. All procedures were approved in advanced by the local Animal Welfare Ethical Review Body and performed in accordance with the Animals (Scientific Procedures) Act 1986 (amended in 2012).

### Reversed Unilateral Ureteric Obstruction (RUUO)

The R-UUO model was performed as previously described and performed by a surgically trained animal technician (41). Briefly, 8-week-old male C57bl/6JOlaHsd mice (Envigo, Indianapolis, USA) underwent laparotomy and the left ureter was isolated and ligated twice with 6/O black braided silk suture close to the bladder. In mice that required reversible ureteric obstruction, a silastic tube was placed around the ureter immediately proximal to the ligature to prevent excessive dilatation. Following 7 days of obstruction, the ureter was re-anastomosed into the bladder and the peritoneum and skin sutured closed. For single cell experiments, mice were culled at day 2 or day 7 post-UUO or 7 days following ureteric re-anastomosis after 7 days of obstruction by CO_2_ narcosis and dislocation of the neck. For ABT-263 administration experiments, mice were culled at day 35 following reversal.

### Unilateral ureteric obstruction (UUO)

Surgery was performed as above, but without the placement of a silastic tube or reattachment of the ureter.

### Ischemia reperfusion injury (IRI)

Surgery was performed by as previously described.^239 A^naesthesi^a^ was induced with 2% iso-fluorane. Buprenorphine analgesia was administered subcutaneously. A posterior flank incision was made and the left renal pedicle identified and clipped using an atraumatic clamp for 15 min. During the ischemic period, body temperature was maintained at 37^0^C using a heating blanket with homeostatic control (Harvard Apparatus, Boston MA) via a rectal temperature probe. The clamp was then removed, the peritoneum closed with 5/0 suture and the skin closed with clips. The other (right side) renal pedicle was not clamped, and the right kidney left completely intact. One ml sterile saline was administered subcutaneously after surgery. These animals were maintained for 28 days post-IRI before tissue harvest.

### ABT-263

ABT-263 was administered as a dose of 50 mg/kg by gavage daily for 7 days, followed by a break of two weeks, then finally another 7 days. ABT-263 was reconstituted with 10% EtOH,30% PEG 400 (Polyethylene glycol) and 60% Phosal 50 PG. The vehicle was the above mixture without the drug.

### LOC14

LOC14 was orally administered by gavage at 20 mg/kg, once daily following UUO. The drug was reconstituted, as per manufacturers guidelines, by adding each solvent one by one: 10% DMSO, 40% PEG300, 5% Tween-80 an 45% saline.

### Immunohistochemistry

Immunohistochemistry was carried out on-board Bond Rx using Leica Bond Intense R staining kit with protocol ImmPRESS Rat using a Bond rx Autostainer (Leica, Wetzlar, Germany). Sections were first warmed to 60⁰C for a 1 hour then dewaxed in Bond dewax (Leica, AR9222) for 30 mins before antigen retrieval in ER2(Leica, AR9640) at 100^0^C for 20 mins. Endogenous peroxidase was blocked for 5 mins before incubation with 2.5% Normal Goat Serum (ImmPRESS Rat kit – Vector Laboratories, San Francisco Bay Area, USA) TRU for 20 mins. Slides were then exposed to P21 1/150 ab for 35 mins (Hugo291, Ab107099, Abcam, Cambridge, UK.). Slides were then immersed in ImmPRESS Polymer Reagent RTU for 30 mins, mixed DAB intense for 5 mins and counterstained in Haematoxylin before dehydration, clearing and mounting. Slides were imaged on the brightfield microscope Zeiss Axioskop (Zeiss, Oberkochen, Germany) where 5 random fields were imaged from each organ’s cortex. Tubules were considered positive if they contained any cells with positive nuclear p21 staining.

### Picrosirius Red staining

Formalin-fixed paraffin-embedded slides were immersed twice in xylene for 5 minutes to dewax. These were then moved through graded ethanol concentrations (100%, 75%, 65%) for 5 minutes each, followed by deionised water for 2 minutes, followed by tap water for 10 minutes. Slides where then lowered into 0.5% Direct Red in saturated picric acid solution (239801, Sigma Aldrich) for 2-4 hours. Next slides were rinsed twice with methylated spirit (100% IMS), rapidly dehydrated in 3 changes of 100% ethanol for 20 seconds each (with agitation) before clearing in xylene and mounting.

Whole slide images were acquired using the brightfield capabilities of an Axioscan Z1 slide scanner (Zeiss). The percentage of Sirius red staining per cortex per organ was then quantified in ImageJ version 1.52p. This was performed by importing .czi files, where each organ formed a separate region of interest. The cortex from each organ was selected using the polygon tool, taking care to exclude any major blood vessels, renal capsule that was incompletely removed during preparation or other obvious artefacts. Following conversion to an RGB stack the thresholding tool was used to “gate in” any Sirius red staining areas, using the coloured images as a guide. This threshold was then kept identical for all images and the area of red stain gated in was expressed as a percentage the selected cortex.

### Sa-B-gal

Frozen sections were cut, warmed to room temperature and air dried for 20 mins before fixing with 500 μL of 4% PFA for 5 mins. These were then rinsed with PBS-MgCl_2_ at PH 5.5 before incubation with fresh SA-β-Gal staining solution - 1 mg/mL X-Gal (Teknova, Hollister, CA 95023, U.S.A) in dimethylformamide, 40 mM citric acid/sodium phosphate (pH 6.0), 5 mM potassium ferrocyanide, 150 mM NaCl, and 2 mM MgCl2at ambient CO_2_ at 37.4°C for 14 hours. Slides were then rinsed and counterstained with nucleofast red (Sigma) before mounting.

Whole slide images were acquired using the brightfield capabilities of an Axioscan Z1 slides scanner (Zeiss). Following a similar method to the Sirius red workflow above, thresholding in Image J allowed quantification of positive staining as a percentage of total renal cortex.

### Cystatin C measurement

Cystatin C ELISA for Kidney Function: Tail vein blood was recovered from mice in an equal volume of 4% Citrate buffer before centrifugation at 5000g for 5 minutes at 4°C. 2ul of the clear phase was transferred into 250ul PBS -/- with 0.5% BSA. The mouse cystatin C ELISA (R&D; Duoset ELISA kit) was carried out according to the manufacturer’s instructions.

### RNA extraction for qPCR

Total RNA from cortical kidney tissue was isolated using the RNeasy kit (Qiagen, Hilden, Germany) following the manufacturer’s instructions. For qPCR analysis of targeted gene expression, cDNA was synthesised from 1µg of template RNA using the QuantiTect Reverse Transcription Kit (Qiagen). VWR Perfecta qPCR mastermix with TAQman (Thermofisher) gene expression assays on a Roche Lightcycler 480 (Roche, Basel, Switzerland) using standard protocol. For human RPTEC cells, the TAQman gene expression assays used were *MDK* (Hs00171064_m1), *LAMA3* (Hs00165042_m1), *PROS1* (Hs00165590_m1), *NRP1* (Hs00826128_m1), *EGFR* (Hs01076090_m1), *JAG1* (Hs01070032_m1), *EPHB2* (Hs00362096_m1), *PDIA3* (Hs00607126_m1), *MAL2* (Hs00294541_m1), *LGALS3* (Hs00173587_m1). Expression levels were normalized for GAPDH (Hs02758991_g1), *HPRT1*(Hs02800695) and *PPIA* (Hs04194521) averaged expression and presented as fold increases over day 7 non-irradiated cells analysed in parallel. For murine kidneys, the following TAQman gene expression assays were used: Cdkn2a (Mm00494449_m1), Cdkn1a (Mm04205640_g1), Il6 (Mm00446190_m1) and Il10 (Mm01288386_m1). mRNA expression levels were normalized for HPRT (Mm03024075_m1) expression and presented as fold increases as compared to uninjured animals. A relative quantification approach was taken using the ΔΔCT-Method. The lightcycler 480 software was used to calculate the Cp values of the target and the reference genes based on an assumption that the efficiency = 2. Data was then presented as fold change compared to uninjured control animals.

### Bulk RNA-Seq

Prior to RNA sequencing, RNA integrity was checked using Genechip (Thermofisher), with only samples with RIN >7 used to generate libraries for subsequent analysis. A PolyA library was constructed and run on an Illumina Novaseq using 5×150bp paired-end sequencing at a depth of 50M reads per library. FastQC was used for initial quality control assessment before trimming was performed with Cutadapt. This followed the standard procedure of first trimming low quality base calls from the 3’ end of the read, before adapter trimming using the default settings-using the first 13 bp of Illumina standard adapters (’AGATCGGAAGAGC’). Reads were then mapped to GRCm38.68 mouse reference transcriptome, mm10 (Ensembl 93) using pseudo-aligner Salmon. A quantification table of the transcripts is generated from this step for subsequent differential expression analysis. Mean % of transcripts aligned across all samples was 76%. A summary of the pseudoalignment statistics can be found in supplemental table 4. Differential expression analysis was performed using DESeq2.(42) This workflow imports the transcript-level quantification data output by Salmon, aggregating to the gene-level with *tximeta*.(43) The starting object had 25719 unique features, 2034 were removed due to minimal expression, leaving 23685 features. Initial QC was performed using PCA plots data having applied a variance stabilizing transformation, with the “blind” argument set to “True”. (This is later set to “False” for downstream Analysis.) All statistical analyses were performed on raw counts, as required by DESeq2, and exploratory visualisations such as covariance matrixes and PCA were performed on transformed data following the variance stabilizing Transformation (VST) implemented in DESeq2. This function estimates this dispersion trend by sub-setting on a small number of genes chosen deterministically, to span the range of genes’ mean normalized count. DESeq2 feature counts and other functions were run on the tables of counts to determine differentially expressed genes between groups (uninjured day 0, day 7 post R-UUO (early repair) and Day 42 post ABT-treatment/matched vehicle group (late -repair). Results were considered statistically significant at an adjusted p < 0.05. in R.

### Data availability

Data have been deposited in the National Center for Biotechnology Information Gene Expression Omnibus database accession # GSE157866 (reviewer token: yhufmocgvhcfjsj) and GSE140023.

## Single cell workflow and bioinformatic methods are found in the supplemental documentation

### Cell culture - Human renal peripheral tubular epithelial cells

Work with human renal peripheral tubular epithelial cells (hTPTECs) were maintained in DMEM-F-12 + Glutamax-1 supplemented with hTERT immortalised RPTEC growth kit (ATCC) and 50mg/ mL Gentamicin. Cells were grown at 37°C in 5% CO_2_. HRPTEC cells were plated at 1 x 10^6^ cells per well of a 6 well plastic culture dish. After overnight culture, these cells were exposed to 10 Gy radiation. Cells were incubated for 7 days with media change every 3-4 days. To induced senescence 100,000 PTECs were seeded onto a 12-well plate. The following day, they were exposed to 10Gy Gamma radiation. Cells were analysed on day 7 and day 14 alongside non-irradiated control groups.

### Cell culture - Human renal fibroblasts

Human primary kidney Fibroblasts (Cell biologics, Chicago, Illinois, United States) were plated at 3 x 10^6^ cells per well of a 6 well plastic culture dish plates pre-coated with gelatin based coating solution (Cell biologics). Cells were maintained in fibroblast medium with added foetal calf serum (Cell biologics). Cells were grown at 37°C in 5% CO_2_. 24 hours before treated with protein of interest, media was swapped for serum free medium to ensure a “serum starved” phenotype to minimise baseline activation of cells. Cells were then treated with PDIA3 (10ng/ml) +- TGFB (10ng/ml). 72 hours later cells were collected for qPCR.

### siPDIA3 down- and pPDIA3 up-regulation

For siRNA-mediated gene knockdown, 4×10^6^ cells were transfected with siRNA oligonucleotide specific for the knockdown of PDIA3 expression in human cells (Sense strand: 5’ ACCTCGTCCTTCACATCTCACTAACATCAAGAGTGTTAGTGAGATGTGAAGGACTT 3’), (Antisense strand: 3’ CAAAAAGTCCTTCACATCTCACTAACACTCTTGATGTTAGTGAGATGTGAAGGACG- 5’) were designed in our laboratory and synthesised by Eurofins MWG Operon (Germany) as control for knock-down efficiency non-specific controls from Eurofins MWG were used. TK188 cells with 70% confluence were transfected with the siPDIA3ERP57 using transfection reagent, Lipofectamine 2000 (Invitrogen) according to the manufacturer’s protocol. Transfection reagent was removed after 6h replaced with normal complete culture medium.

For the overexpression of PDIA3, 70% confluent TK188 cells were transfected either with pPDIA3 (pcDNA3.1-ERP57, a gift from Dr. Neil Bulleid, University of Glasgow) using transfection reagent, LTX reagent with PLUS reagent (Invitrogen) according to the manufacturer’s protocol. After 6 h of transfection media was replaced with normal complete media supplemented with 0.5 mg/ml G-418 (Invitrogen) as a selection factor for stable transfection.

Both overexpression and knockdown of PDIA3 was confirmed by performing western blotting using mouse anti-PDIA3 antibody (Enzo Life Sciences, Lörrach, Germany)

The activation of TGFß1 pathway in the siPDIA3 and pPDIA3 treated TK188 was investigated by monitoring the phosphorylation of SMAD2 using the rabbit-anti-pSmad2 antibody phosphospecific to Ser465/467 (Merck Millipore Billerica, MA, USA).

### Measurement of cell viability

6×10^3^ TK188-cells/well was seeded in 96-well plated and treated with either siPDIA3 or pPDIA3 for 24 h. Medium was removed, and after washing in phosphate-buffered saline (PBS) the cells were incubated for further 24 h in 10 ml serum-free DMEM. Purified human *TGFβ1* (5 ng/ml) (R&D systems, Minneapolis, MN, USA), was added to the medium and the cells were incubated for additional 48 h. Cell viability was assessed using cell Proliferation Kit I (MTT), a colorimetric assay for the non-radioactive quantification of cell proliferation and viability (Roche Applied Bioscience, Mannheim, Germany).

For the trypan blue protocol to determine the impact of the different treatments on cell viability, the fibroblast cell line TK188 were cultured in 6 well plate and treated for 48 h as follow: no treatment (control), PDIA3, TGFB1, PDIA3 & TGFB1, PDIA3 & Loc14, TGFB1 & Loc14, or TGFB1 & PDIA3 & Loc14. The cells were then suspended in PBS and mixed to equal volume (100 µl each) with the 0.4% trypan blue staining solution. To determine the percent of viable cells, 10 μL of the stained cell suspension were counted on a Countess II FL Automated Cell Counter (thermo Scientific).

To investigate whether ERp57 is involved in the TGFß-1 pathway regulation, TK188 cells were treated with human recombinant ERp57 (Novus Biologicals Wiesbaden, Germany) with and without TGFß-1 and/or Loc14. And the activation of TGFß-1 pathway was monitored by investigation the expression of the pSMAD and Smad anchor for receptor activation (SARA) using rabbit-anti-pSmad2 antibody phosphospecific to Ser465/467 (Merck Millipore Billerica, MA, USA), and the rabbit-anti-SARA antibody (Cell Signaling, Frankfurt am Main, Germany).

### Measurement of free active TGF- β1

For measuring free active TGF- β1 in media exposed to hRPTECs, 80,000 cells were seeded into 24 well plates. For 48 hours, they were maintained at 37°C in 5% CO2 in in-house media composed of DMEM:12 + Glutamax-1 supplemented with the following; Geneticin (Life Technologies) 50mg in 1 ml, Hydrocortisone (Merck) 12.5ug, Ascorbic Acid (Sigma-Aldrich) 1.75mg, Insulin-Transferrin-Selenium-Ethanolamine (ITS-X) (Thermofisher Scientific) 10mls, Triiodo-L-Thyronine (Merck) 6pM, Prostaglandin E1 (Merck) 12.5ug, recombinant human Epidermal Growth Factor (Promega) 5ug and HEPES (Sigma-Aldrich) 1.13g in 4mls of 1N Sodium Hydroxide.

The cells were then maintained for a further 24 hours in serum starved media composed of DMEM:12 + Glutamax-1 supplemented only with fetal bovine serum 0.1% and penicillin-streptomycin. L-TGFb1 (299-LT-005, R&D systems) was then incubated at 5000pg/mL for 1 hour at 37°C at 5% CO_2_, in the presence or absence of PDIA3 at 2µg/mL (ab92937-100ug, Abcam), with or without LOC14, a PDIA3 inhibitor, used at 100µM.

LEGEND MAX™ Free Active TGF-β1 ELISA Kit (BioLegend) was used to measure active TGF-b per manufacturer’s instructions. The cytokine concentration in each supernatant was extrapolated from a standard curve.

### Power and statistical analysis

Animal group size was determined from previous pilot experiments. Normality was assessed with a Shapiro-Wilk test and visualization of data distributions. Levene’s test was used to test for homogeneity of variance across groups. Comparisons between two unpaired, normally distributed data points were carried out via a T-test and between two unpaired, non-normally distributed data points were carried out via Mann-Whitney test. Comparisons between multiple groups were performed with one-way ANOVA with Tukey’s multiple comparison test if parametric or Kruskal-Wallis test if a nonparametric distribution. All statistical analysis was performed using R version 4.

## Author Contributions

EOS and DF conceived and designed the study. EOS, KM performed flow cytometry sorting experiments for library preparation, KM performed hPTEC work and animal experiments alongside DF, EOS. EOS and CCarvalho performed fibroblast experiments. DB and CCarvalho performed cell culture and measurement of free TGFb. MD, EOS produced immunofluorescence images. EOS, CCairns prepared cDNA libraries, and CCairns supported tissue staining and qPCR. KG and AL provided human biopsy samples. NH provided single cell platform, resources and expertise. KK supervised single cell RNA-Seq workflow and analysis. TC, BC and LD provided intellectual input and designed single cell RNA-Seq experiment and supervised analysis. RC performed staining, qPCR and analysis of same. RB, LD performed R-UUO surgeries. HD performed animal experiment and confirmed independently the EOS & DF results, HD, GHD performed fibroblast experiments and WB. HD, MZ, provided intellectual input and guidance. EOS performed statistical, microscopic and bioinformatic analysis of data, generated figures and wrote manuscript. DF performed animal surgeries. DF and JH supervised the project and provided intellectual input and guidance and manuscript editing. All authors critically reviewed and approved the manuscript before submission.

## Acknowledgements

EOS is funded by Kidney Research UK clinical training fellowship (TF_006_20161125). DAF was supported by an Intermediate Fellowship from the Wellcome Trust (100171/Z/12/Z, The effect of aging upon renal injury and repair).

We would like to thank J R Wilson-Kanamori for his invaluable insight into single cell analysis.

