## supplemental figure for "Single cell analysis of senescent epithelia reveals targetable mechanisms promoting fibrosis"

### Supplemental Figure 1

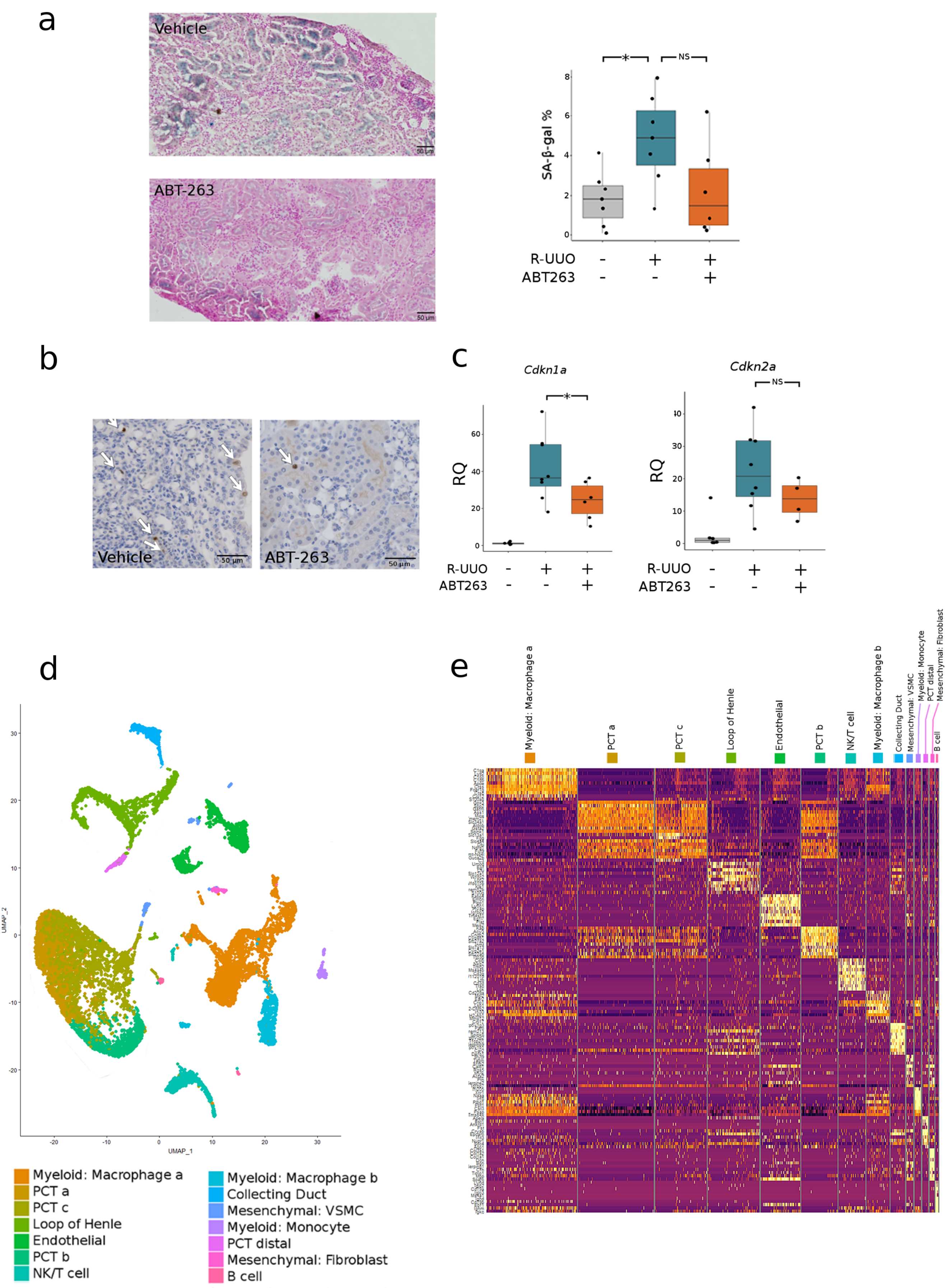

**Supplemental Figure 1**

**Senescent cells are persistent following renal injury, incomplete repair.**

a) Quantification of percentage SA-β-gal per whole kidney. * denotes P val <0.05. Vehicle treated kidneys 4.8% (s.d. 2.3) vs ABT-263 treated 2.2 (s.d. 2.2), two-sided t-test, *P* =0.07, CI -5.3-0.2.

b) Representative images of P21 immunohistochemical staining.

c) qPCR data showing *Cdkn1a* and *Cdkn2a* reduction post ABT-263 treatment.

**Classification of cells in single cell RNA-Seq data**

d) UMAP of 17136 cells from libraries pooled from mice that underwent sham, UUO-2, UUO-7 or R-UUO (n=3/time-point) classified by a cell cluster.

e) Heatmap of top 10 genes per cluster by fold change, calculated using Wilcoxon signed-rank test. The colour scheme is based on z-score distribution.

### Supplemental Figure 2

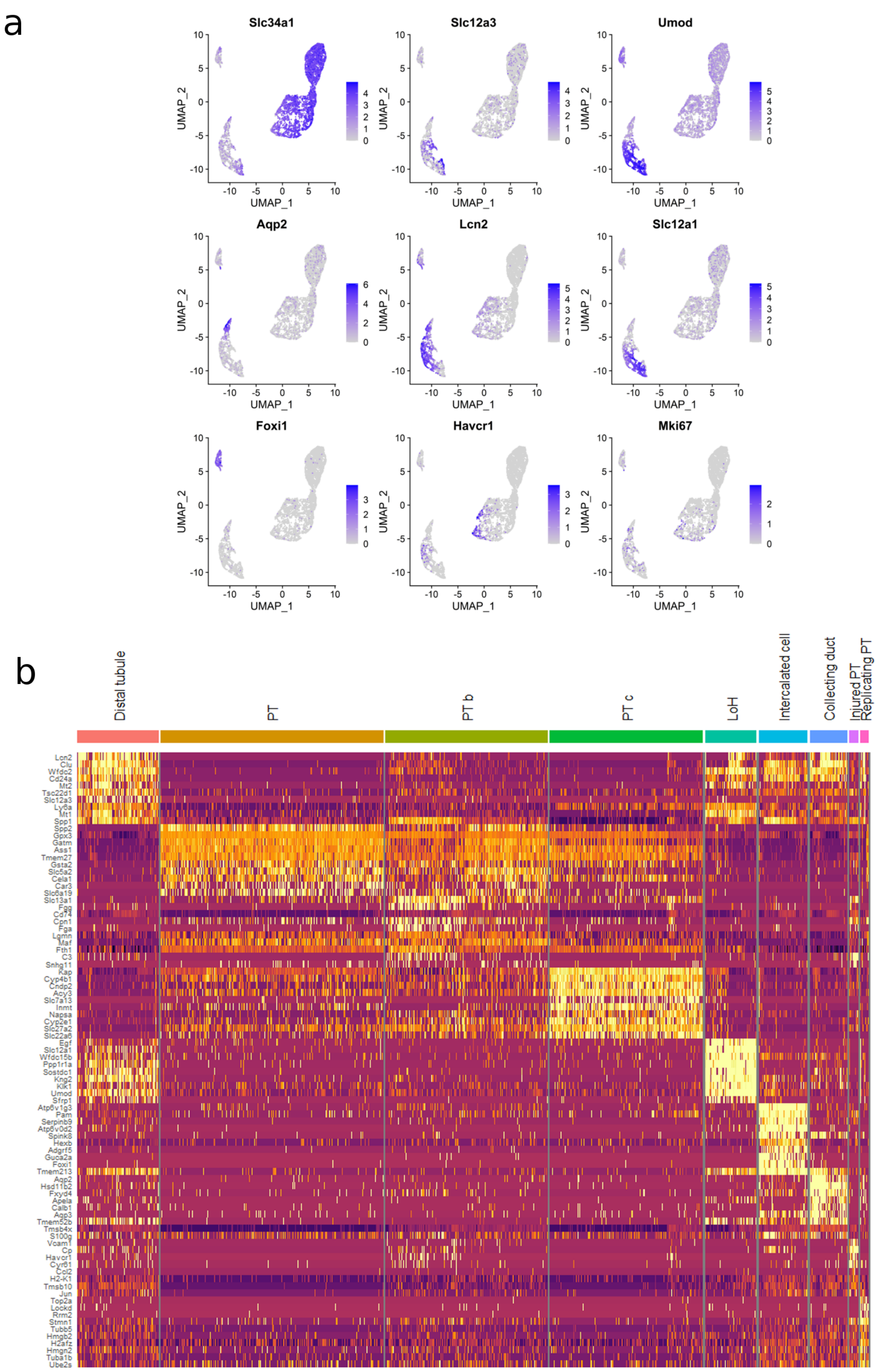

**Supplemental Figure 2**

**Epithelial cell markers in single cell RNA-Seq**

a) Classical epithelial cell gene expression projected onto UMAP. Colour intensity proportional to expression.

b) Heatmap of top 10 genes per epithelial cluster by fold change, calculated using Wilcoxon signed-rank test. The colour scheme is based on z-score distribution.

### Supplemental Figure 3

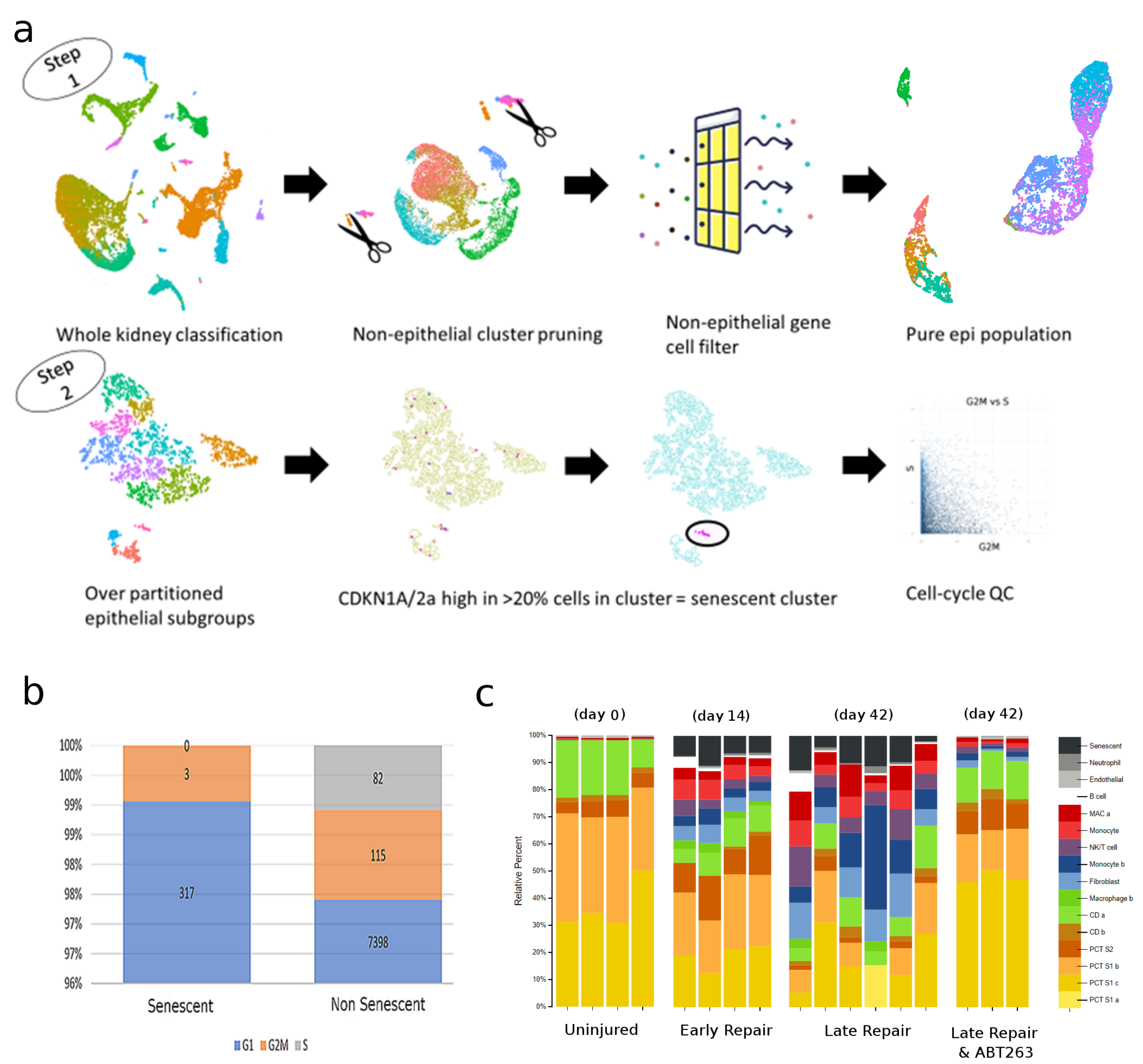

**Supplemental Figure 3**

**Identification of senescent cells in single cell RNA-Seq data**

a) Schematic depiction of the workflow to identify senescent cells post-hoc, as described in detail in methods.

b) Proportions of cell phase assignments.

c) To confirm our single cell derived signature was appropriately detecting senescent cells, we used cibersortx (see methods) to deconvolute our previous bulk RNA-Seq data into cell type proportions. Here we see incomplete recovery post injury associated with senescence cells, as well as myeloid cell expansion and fibroblast activation, and a depletion of senescent cells with ABT-263 administration.

### Supplemental Figure 4

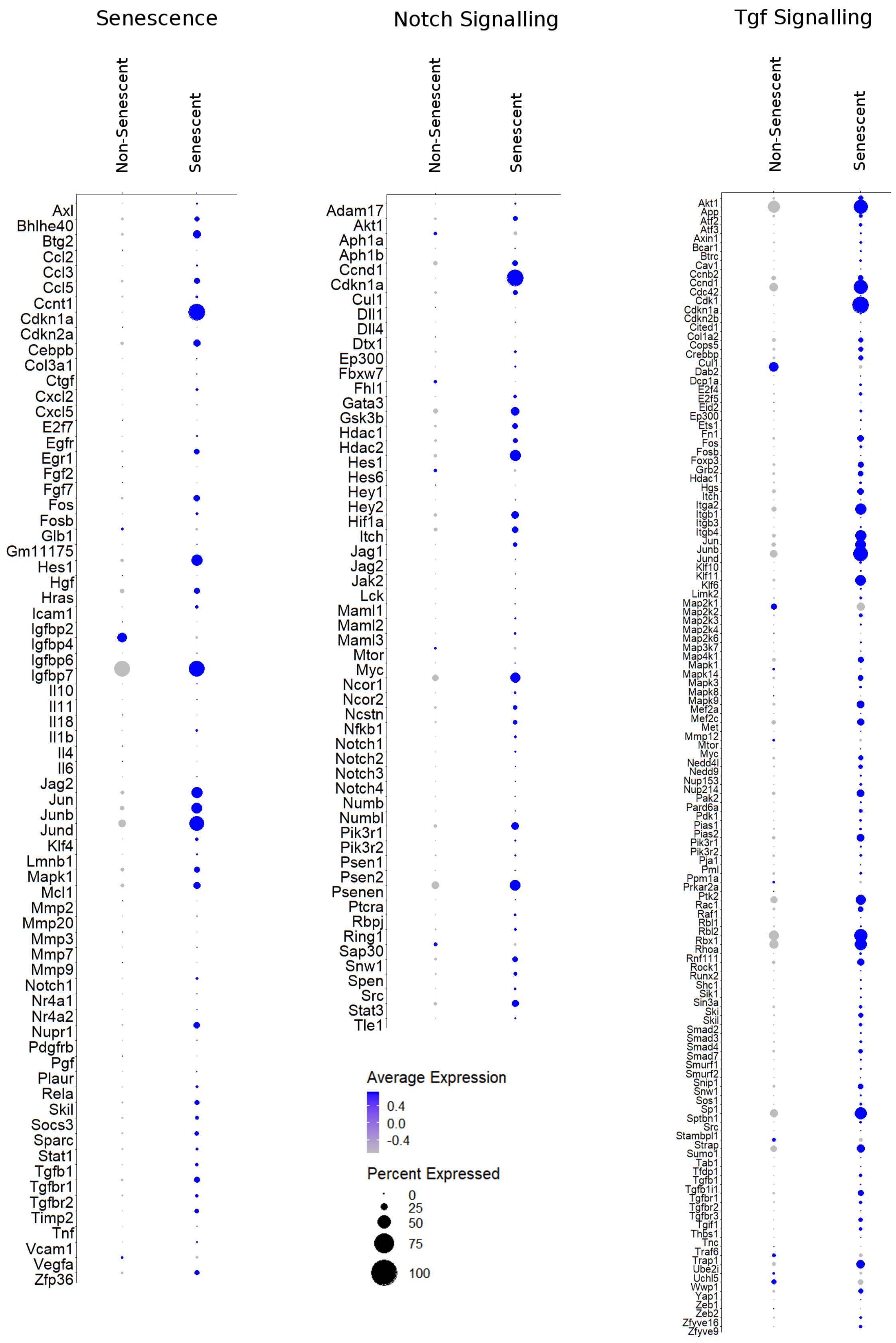

**Supplemental Figure 4**

**Senescent cells express Tgf and Notch pathways**

a) Senescent cells express genes pathways typical of senescence, as well as the Notch and Tgf signalling pathway. Size of dot represents the percentage of cells in each cluster expressing the gene, colour indicates average gene expression per treatment.

### Supplemental figure 5

#### Ischemia Reperfusion Injury (GSE 140010)

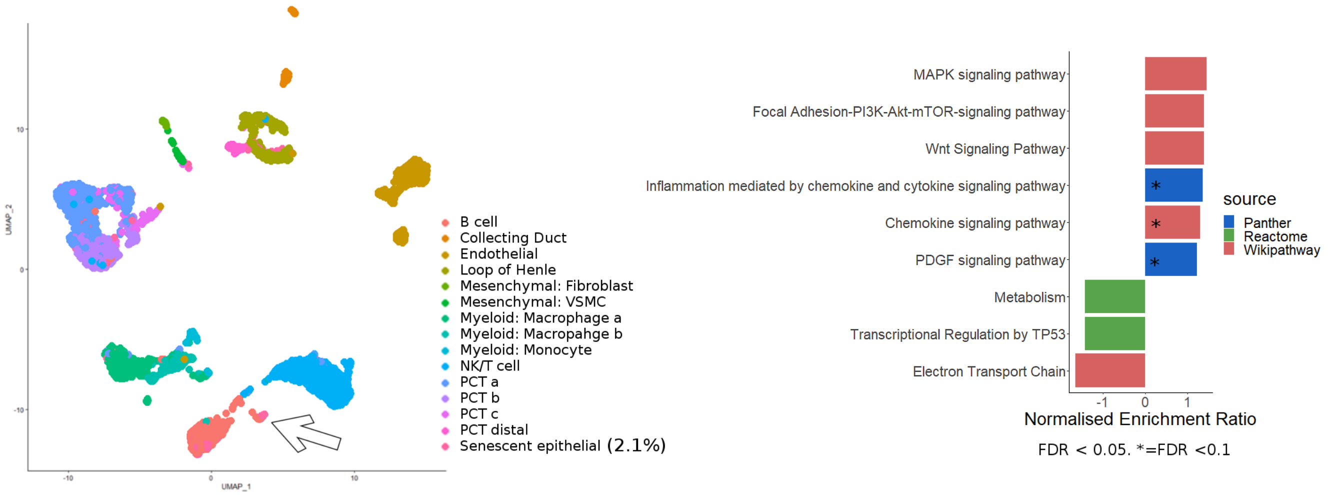


#### Ischemia Reperfusion Injury (GSE 139506)

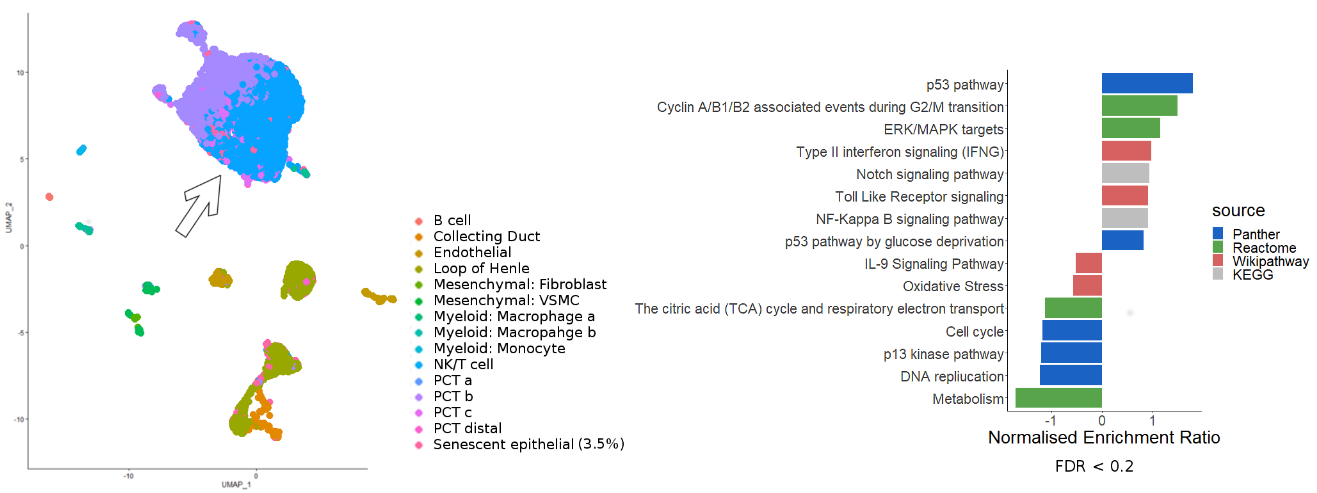


#### Tabula Muris Senis (GSE132042)

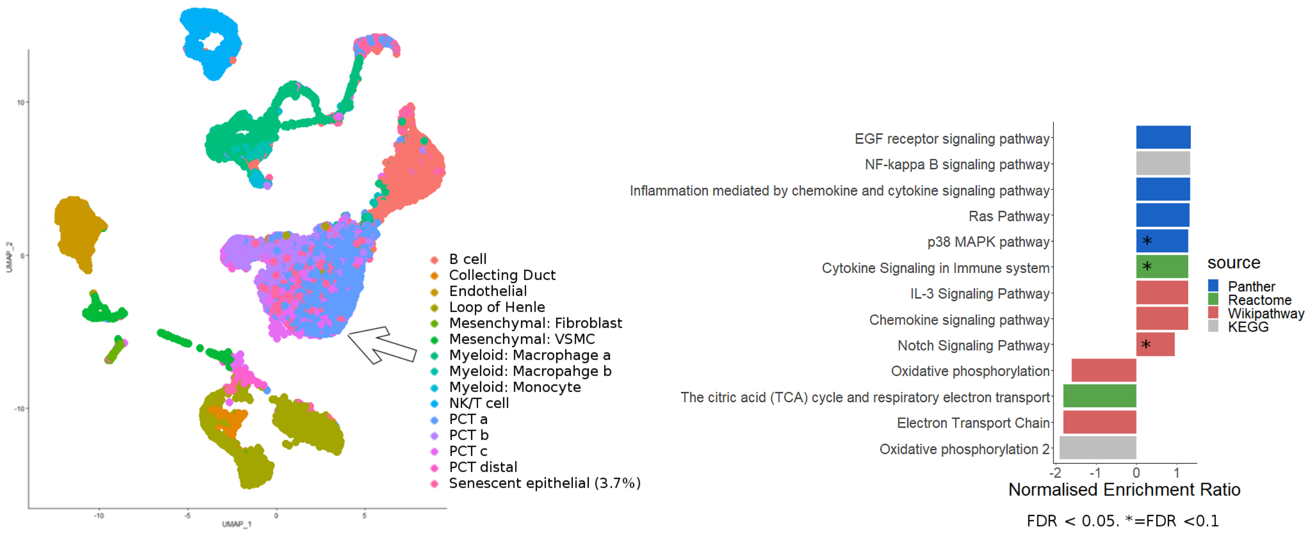

**Supplemental Figure 5**

**Single cell classification validation in publicly available datasets**

Senescent cells were classified in publicly available datasets - two ischemia reperfusion injury models and tabula muris senis, which includes aged mice. GSEA was performed on the DEG between the senescent epithelial cells as compared to non-senescent PCT. Positively enriched pathways are higher in senescent epithelia relative to non-senescent.

### Supplemental Figure 6

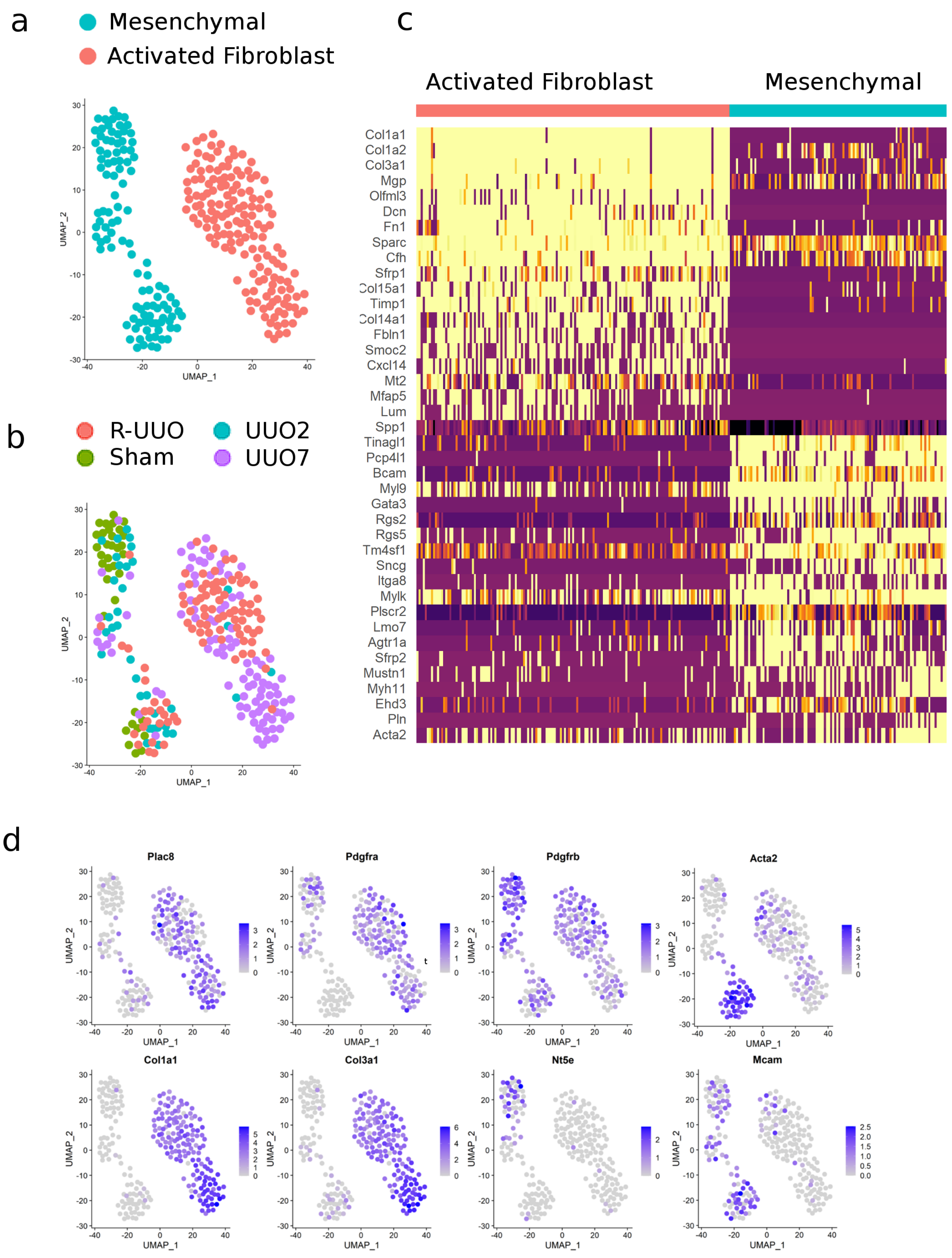

**Supplemental Figure 6**

**Single cell fibroblast classification**

a) UMAP of 259 mesenchymal cells, coloured by classification.

b) UMAP of 259 mesenchymal cells, Coloured by time along injury.

c) Top 40 DEG between activated fibroblasts and mesenchymal cells.

d) Gene expression of key mesenchymal genes showing typical expression of activated genes in activated myofibroblasts, allowing classification of a distinct cluster as myofibroblasts. Colour intensity proportional to expression.

Supplemental figure 7

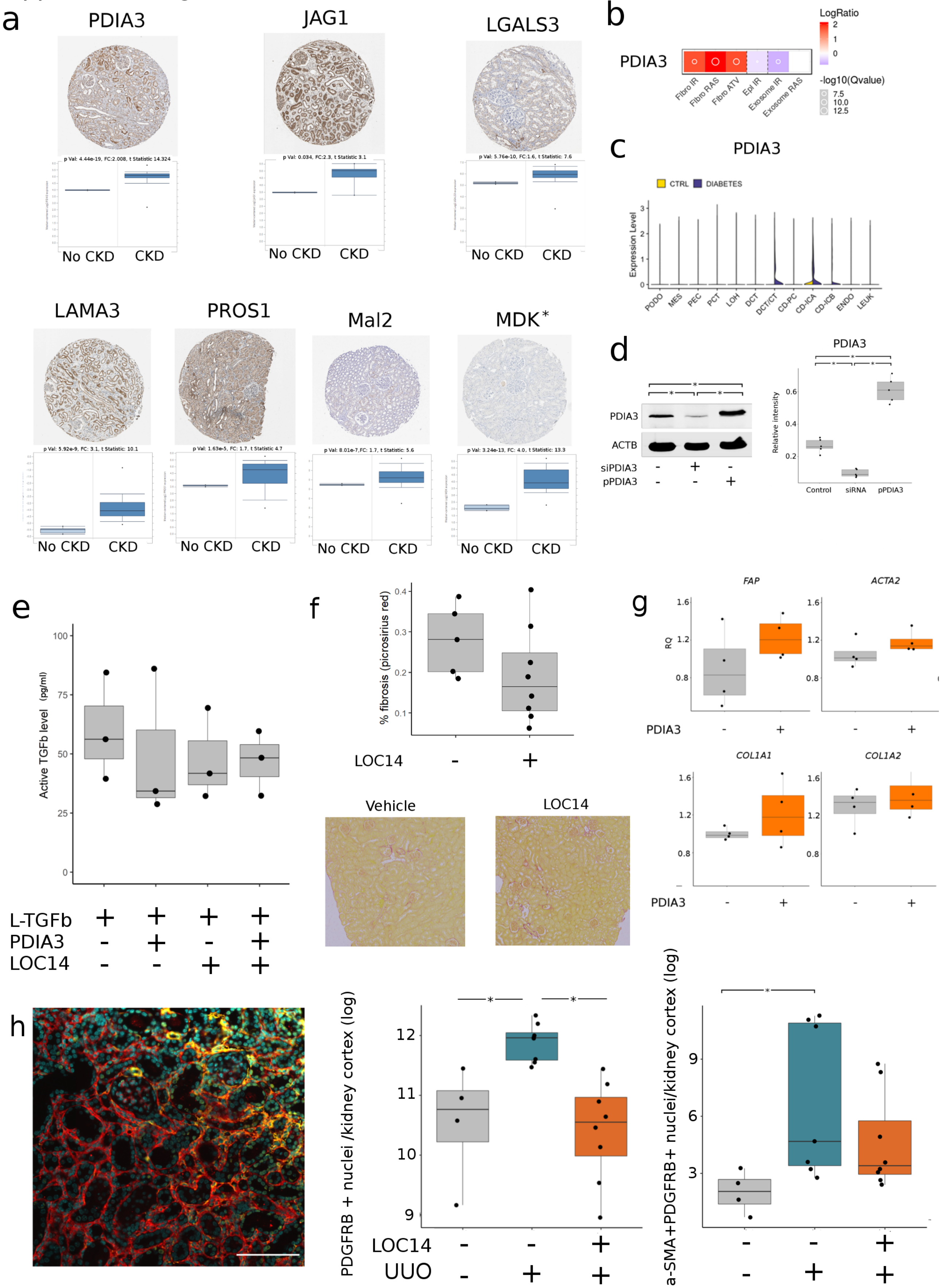

**Supplemental Figure 7**

**Target molecules are expressed in human kidneys and enriched in CKD**

a) Staining of human protein atlas samples (ages 26-70) demonstrates all candidate molecules can be found in the tubular epithelial *(except MDK). The Nakagawa dataset on nephroseq (www.nephroseq.org) shows each gene to be significantly increased in CKD as compared to patients without CKD.

b) Heatmap showing presence of the PDIA3 in the SASP atlas. Expression relative to control. Fibro=fibroblast. Epi=renal tubular epithelial. IR= irradiation, RAS=RAS overexpression, ATV=Atazanavir. Exosomes are from irradiated fibroblasts. Numeric Value is log2 ratio between senescent and non-senescent cells. Source www.saspatlas.com

c) There is increased *PDIA3* expression in human tubular cells in patients with diabetic nephropathy as compared to those without. Source: ttp://humphreyslab.com/SingleCell/ data: Wilson et al

**Western Blot & quantification**

d) Western Blot demonstrating that a Small interfering RNA (siRNA) directed against PDIA3 reduced PDIA3 synthesis in TK188 fibroblast cells, and that a PDIA3 plasmid (pPDIA3) resulted in higher PDIA3 synthesis. Quantification of PDIA3 from below blot.. Relative intensity is the densitometric value of the western blot bands divided by the intensity of the corresponding Actin band. Comparisons tested using one way ANOVA with Bonferroni correction. Significance set at P adj< 0.05

**PDIA3 Does not increase free active TGFb**

e) Quantification of free TGFb using commercially available ELISA kit. No statistical difference between groups, suggesting PDIA3 does not increase active TGFb *in vitro.*

**Loc14 has no effect on kidney fibrosis**

f) Uninjured kidneys treated with Loc14 or vehicle. Loc14 treated 0.19% (s.d. 0.11) vs 0.27% (s.d. 0.08), t-test, P =0.15, CI -0.03-0.2

**PDIA3 has no effect on fibroblast activation in isolation**

g) qPCR of human kidney fibroblasts treated with PDIA3. No significant difference measured by two-sample Wilcoxon test.

**Loc14 reduced fibroblast number post UUO**

h) Representative image of dual + staining for PDFRB and a-SMA. Scale bar = 100 microns. DAPI+ nuclei in cyan, PDGFRB=red, a-SMA=Yellow.

Number of nuclei in positive cells across entire kidney cortex in mice during UUO injury with and without Loc14 treatment was quantified and normalised to the area of cortex selected. This result was then logged (natural log).

The centre line represents the mean, the box limits the first and third quartiles, the whiskers +- 1.5 * IQR and the points all the data

Supplemental figure 8

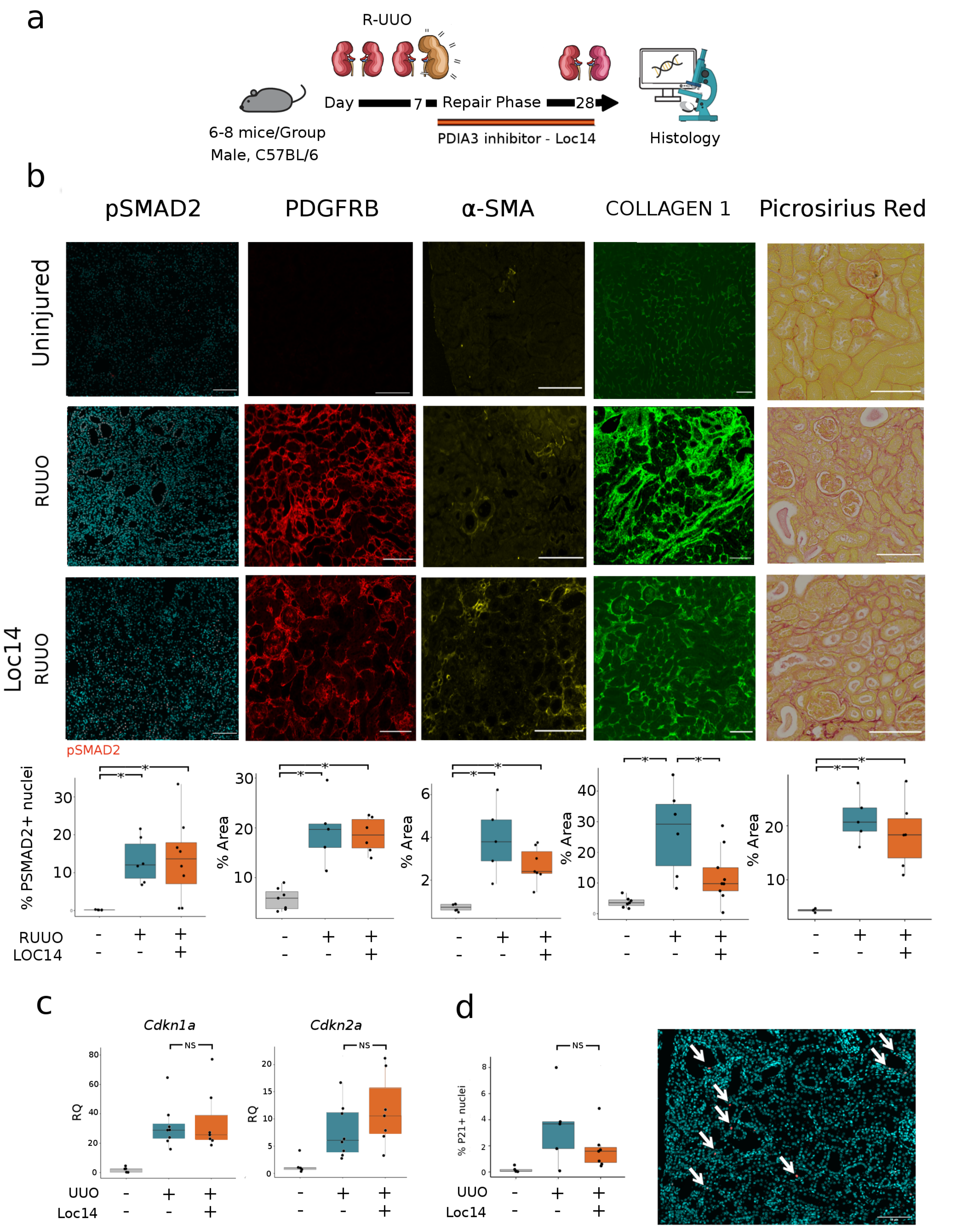

Supplemental Figure 8

Loc14 has no effect on number of senescent cells

a) experiment schematic showing Loc14 administration to mice post R-UUO surgery

b) Representative images and quantification immunofluorescence and of picrosirius red staining of kidneys. Quantification of total staining per renal cortex. * denotes P val <0.05.

PSMAD2 uninjured 0.172% (s.d. 0.6). R-UUO & Vehicle treated 13.2% (s.d. 6.07) vs R-UUO & Loc14 treated 13.7% (s.d. 10.9). ANOVA, *P*=0.935 95% CI: -7.66-5.8

Pdgfrb: uninjured 5.69% (s.d. 2.23). R-UUO & Vehicle treated 19.5% (s.d. 6.78) vs R-UUO & Loc14 treated 18.6% (s.d. 3.57). ANOVA, *P*=0.99 95% CI: -10.6-11.61

a-SMA: uninjured 0.72% (s.d. 0.16). R-UUO & Vehicle treated 3.91% (s.d. 1.7) vs R-UUO & Loc14 treated 2.7% (s.d. 0.82). ANOVA, *P*=0.142 95% CI: --2.77-0.34

Collagen 1: uninjured 3.75% (s.d. 1.73). R-UUO & Vehicle treated 26.8% (s.d. 14.4) vs R-UUO & Loc14 treated 12.4% (s.d. 8.8). ANOVA, *P*=0.0.22 95% CI: -26.98-1.

Picrosirius Red: control & vehicle treated 4.3% (s.d. 0.32). R-UUO & Vehicle treated 21.3% (s.d. 4.4) vs R-UUO & Loc14 treated 18.4% (s.d. 6.3) a difference of 2.9%, (C.I -4.9-10.8) P=<0.59

For boxplots, the centre line represents the mean, the box limits the first and third quartiles, the whiskers +- 1.5 * IQR and the points all the data. Scale bar = 100 microns.

c) qPCR of whole kidney samples, NS= no significant difference between groups.

Cdkn2a: 11.5 (s.d. 6.6) vs 7.7 (s.d. 4.9), t-test, P =0.2 CI -2-10.5. Cdkn1a: 34.8 (s.d. 21.4) vs 31.7 (s.d. 14.9), Wilcoxon ranked sum test, P =0.8 CI -12-22.

d) Immunofluorescence of P21(ab188224) showing nuclear positivity (white arrows). P21 positive nuclei quantified as percentage of total DAPI positive nuclei. Loc14 treated 3.4% (s.d. 2.9) vs 12.1% (s.d. 11.7), t-test, P =0.079, CI -1.2-18.6.

Kidney digestion for single-cell sequencing.

Immediately after culling, mice were perfused with 10 mL PBS. Kidneys were excised, decapsulated and placed in ice-cold PBS. Equal portions of renal cortex from each mouse were finely minced in digest buffer [4.25mg/mL Collagenase V (Sigma-Aldrich St. Louis, Missouri, USA), 6.25mg/mL Collagenase D (Roche, Basel, Switzerland), 10mg/mL Dispase (Thermo-Scientific, Waltham, Massachusetts, United States) and 300μg/mL DNase (Roche, Basel, Switzerland) in RPMI 1640 (10% fetal calf serum, 1% penicillin/streptomycin/L-glutamine)] prior to homogenisation in gentleMACS C-tubes using the gentleMACS dissociator (Miltenyi Biotec, Auburn, California, USA). Samples were incubated at 37ºC with shaking to maximise digestion. The kidney suspension was then subjected to a second gentleMACS homogenisation and digestion neutralised with an equal volume of FACS buffer (PBS, 2mM EDTA and 2% FCS). Kidney cell suspensions were then passed sequentially through 100 μm, 70 μm and 40 μm sieves. Any residual red blood cells were lysed by RBC lysis buffer (Sigma-Aldrich St. Louis, Missouri, USA). Cells were re-suspended in ice-cold FACS buffer ready for use.

Single-Cell droplet library preparation

For single-cell RNA-Seq analysis on the 10x Genomic platform (10x Genomics, Pleasanton, California, USA), single-cell suspensions from renal cortex were prepared, from pools of 3 animals from each group, as outlined by the 10x Genomics Single Cell 3' v2 Reagent kit user guide. 50,000 live (DAPI-) cells were sorted on the BD FACS ARIA II. Samples were washed twice in PBS followed by centrifugation at 500g for 5 minutes at 4oC. Sample viability was assessed using trypan blue with an automated cell counter (Bio-Rad, Hercules, California, USA) and the appropriate volume for each sample was calculated. The chip was loaded with 10700 cells per lane.

After droplet generation, samples were transferred onto a pre-chilled 96-well plate, heat-sealed and reverse transcription was performed using a c1000 touch thermal cycler (Bio-Rad, Hercules, California, USA). After reverse transcription, cDNA was recovered using the 10x Genomics Recovery Agent and a Silane DynaBead cleanup (ThermoFisher, Waltham, Massachusetts, U.S.A) was performed. Purified cDNA was amplified and cleaned using SPRIselect beads (Beckman, Brea, California, U.S.A). Samples were diluted at 4:1 (elution buffer (Qiagen, Düsseldorf, Germany)/cDNA) and an initial concentration check performed on a Qubit fluorometer (Invitrogen/ThermoFisher, Waltham, Massachusetts, U.S.A) to ensure adequate cDNA concentration. Final cDNA concentration was checked on a bioanalyzer (Invitrogen/ThermoFisher, Waltham, Massachusetts, U.S.A).

Single cell RNA-Seq

We generated libraries (3 biological replicates per library) from 4 time points during the R-UUO procedure. These single cell libraries were generated from whole kidney digests and prepared using a high-throughput droplet-based library preparation workflow (10x v2). These libraries were then pooled and normalised by molarity before being sequenced across 4 lanes on a single illumina flow cell. Sequencing was performed on an Illumina HiSeq platform with a target of ~350M PE reads/lane, giving ~525 M PE reads/sample comprising of 2x150bp PE configuration and 8bp index reads. Data was deposited in the National Center for Biotechnology Information Gene Expression Omnibus database (accession #GSE140023)

Single-cell RNA sequencing analysis

For the droplet-based dataset, the cellranger mkfastq wrapper (Cell Ranger Single Cell software suite 2.1.0, http://10xgenomics.com) de-multiplexed the illumina output BCL files to library specific FASTQ files. Subsequently, alignment was performed using the cellranger count function using STAR aligned 2.5.1b (44) against the Ensembl mouse reference genome version GRCm38.68. Correction and filtering of cell barcode and unique molecular identifiers followed, and the retained barcodes were quantified and used to generate a gene expression matrix. Summary sequencing statistics are provided in supplemental table 6.

For our droplet-based dataset, a standard sequence of filtering, highly variable gene selection, dimensionality reduction and clustering were performed using the scRNA-Seq analysis R package Seurat(v2.3.4).(45) Following alignment and initial pre-processing, we began our R workflow with 15,046 unique genes across 7,073 cells in our Sham group, 16,450 unique genes across 5,088 cells in our UUO-2 group, 17,368 unique genes across 7124 cells in our UUO-7 group and 17,227 unique genes across 6,096 cells in our R-UUO group. There were mean of 1218 features per cell. To exclude low-quality cells in both single-cell experiments, we then filtered cells that expressed fewer than 300 genes per 500 unique molecular identifiers, and to exclude probable doublets, cells with >10,000 unique molecular identifiers and <3,000 genes were removed. This will have removed the majority of injured and apoptotic cells.(46) We used a mitochondrial filter to remove cells in which >50% of genes were mitochondrial, consistent with other renal-specific scRNA-Seq projects.(47, 48) This is a higher filter than has been used in non-renal single-cell analysis but reflects the high mitochondrial content in renal tubular epithelial cells. Any gene not expressed in at least 3 cells was removed.

Normalisation was performed using the Seurat package to reduce biases introduced by technical variation, sequencing depth and capture efficiency. We employed the default global-scaling normalisation method “logNormalize” which normalised gene expression per cell by the total expression and multiplies the result by a scaling factor before log transformation. We then scaled the data and regressed out variation between cells due to the number of unique molecular identifiers and the percentage of mitochondrial genes.

The expression matrix subsequently underwent dimensionality reduction using principal component analysis (PCA) of the highly variable genes within the dataset. Using Seurat’s FindVariableGenes function (and computed using the LogVMR argument) we used log-mean expression values between 0.0125 and 3, and a dispersion cut-off of 0.5 to select genes. PCA was performed using these selected genes and 20 principal components were identified for subsequent analysis in each dataset, selected both visually using the elbow point on the elbowplot and via the Jackstraw method.

Further cluster-based quality control was performed in the droplet data using the density-based spatial clustering algorithm DBscan which was used to identify cells on a tSNE map. We initially set an eps value of 0.5 and removed clusters with fewer than 10 cells. The remaining cells were then clustered again with eps value 1, followed by removing the clusters with fewer than 20 cells. Of note this allowed identification of a cluster characterized by high expression of heat shock genes including Fos, Jun, Atf3. This cluster was removed as it was considered to be an artefact of cell stress due to the experimental protocol as recently described.(49) This procedure removed 158 (3.4%) from a total of 4,540 cells in sham mice, 137 (4.4%) of 3,101 cells in UUO-2 mice, 167 (3%) of 5,563 cells in UUO-7 mice and 83 (1.9%) of 4,308 cells in R-UUO mice.

Clusters were then identified using Seurat's FindClusters function, built using the first 20 principal components and a resolution parameter of 0.5-1.5. The original Louvain modularity optimization algorithm was employed. UMAP was then used for further dimensionality reduction and visualisation, which was run on a reduced dimensional space of the first 20 dimensions.

For all single-cell differential expression tests, we used the Wilcoxon rank-sum test to identify a unique expression profile for each cluster, with differential expression tested between each cluster and all other clusters combined. The FindAllMarkers test as implemented in Seurat returns an "adj_pval" (Bonferroni adjusted p-values) and an "avg_logFC" (average log fold change) for each gene. Genes were ranked in order of average log fold change and visualised using heatmaps.

Epithelial Cell Identification

The process of acquiring a dataset of pure epithelial cells required a combination of cluster-based cell pruning and a gene-based cell filter. The clusters were classified by cell type, using the differentially expressed marker genes generated, and recognised markers based on the single cell literature. All epithelial cell clusters were then selected and subsetted on. These were renormalized and rescaled as described above, and were processed again through the pipeline described above, although without the additional DBscan step. Clustering resolution was lowered for each downstream FindClusters function and again clusters were identified based on differentially expressed genes. Any non-epithelial cell clusters were removed, and the data was once again reprocessed and re-clustered. This entire process was repeated three times in total, using successively smaller resolutions used to generate greater numbers of clusters, in an effort to coerce any non-epithelial cells into forming a distinct cluster as the data became cleaner and cleaner. The expression of 52 key non epithelial genes was then assessed and any cell expressing over a predetermined threshold ( typically a log order higher than all other cells) was removed as follows: Endothelium (Ehd3,Plat, Pecam1), Myeloid: (Adgre1, Ccr2, Lyz2, Ly6c1, Cd14, Cd68, C1qa, C1qb), Stromal: (Pdgfra,Pdgfrb,Plac8,Vim,Acta2), Podocytes: (Nphs1,Nphs2,Wt1,Ptpro,Lamb2,Npr3,Podxl,Itga3,Synpo,Npr1,Axl), Lymphoid: (S100a9,Csf4r,Ptprc,Ltb,Cd4,Cxcr6,Nk7,Igkc,Itgae), Cycling( Mki67)

Senescent Cell Identification

Single Cell RNA-Seq fails to sequence Cdkn2a (even at read depths of up to 100,000 reads per cell in our pilot data). Cdkn1a is another marker of senescence but may be expressed in damaged non-senescent cells. Given the transcriptomic signature of senescent cells should be similar enough to each other to partition into a distinct cluster at higher resolutions we used the cell type hierarchy-based classification function of Monocle v2.8.0 to classify the epithelial cells as senescent.(50, 51) The data, previously scaled, normalised and clustered by Seurat was imported into a monocle celldataset. First size factors were estimated using the median ratio method described by Equation 5 in Anders and Huber before tSNE dimension reduction of the first 10 principal components with a low perplexity setting in the range of 15-20.(52) This resulted in 20-25 distinct clusters per dataset, and any cluster with more than 20% of cells expressing Cdkn1a was deemed senescent. Only one cluster fit this criterion in each dataset. These cells were then reanalysed and compared to their non-senescent counterparts. The priority was a highly specific signature, rather than a highly sensitive one, cells which were transcriptionally ambiguous were removed from some clusters to minimise the risks of over classification with would reduce our differential expression’s ability to detect meaningful genes more than under-classification would. While the statistical analysis was performed using the entire dataset, for the purposes of heatmap visualisations, non-senescent data were downsampled to improve legibility of figures.

Cell Phase Assignment

Cell Phase assignment was then performed using the Cyclone package as a QC step to ensure no erroneous classification of cycling cells due to the role of Cdkn1a/2a in the cell cycle. Cyclone is the classification step of the pair-based prediction method described by Scialdone et al.(53) Briefly this function works by using gene expression data to train a classifier for assigning cell phase. We used a training set of mouse cell-cycle markers available from SCRAN. Using a pair-based prediction model, pairs were identified such that the expression of the first gene in the training data was greater than the second gene in a given phase but less than the second in all other phases. Cyclone then calculates, on an individual cell basis, the proportion such gene pairs where the expression of the first gene was greater than the second new data provided. A high proportion of such a pattern from a particular phase training sets, suggested that the cell was likely to belong that phase. Proportions were converted into scores that account for the size and precision of the proportion estimate. Cells with G1 or G2M scores above 0.5 were assigned to the G1 or G2M phases, respectively. The higher score was used if they both exceed 0.5. Interestingly, while cells could have been assigned to S phase based on the calculated S score, a more reliable approach was to define S phase cells as those with G1 and G2M scores below 0.5. In order to allow comparison between the phase specific proportions, cyclone constructs a distribution of proportions by shuffling the expression values within each cell and recalculating the proportion. The phase score was defined as the lower tail probability at the observed proportion. High scores indicated that the proportion is greater than what is expected by chance if the expression of marker genes were independent.

#### Ligand-pair interactions

Heatmaps and dotplots of number of ligand-pair interactions were generated using the CellphoneDB tool (https://www.cellphonedb.org/) developed by the Teichmann Lab, Wellcome Sanger Institute, Cambridge, UK.(54) The lower cut-off for expression proportion of any ligand or receptor in a given cell type was set to 10%, and the number of permutations was set to 1000. The clusters were not subsampled.

#### Pathway Analysis

Gene set enrichment analysis (GSEA) and over representation analysis (ORA) were used to identify enriched pathways based on the differential expressed genes using WebGestalt. For GSEA, we generated a rank for each gene in the list of DE genes. For the GSEA method, categories were discarded if they contained less than 15 genes or more than 500 genes. The Benjamini-Hochberg method was used to correct for multiple testing during ORA, and the top ten enriched categories as ranked by false discovery rate were selected. The reference gene list used was the Illumina mouseref 8. We used pathway gene sets from the protein analysis through evolutionary relationships, PANTHER, Reactome and Kyoto encyclopaedia of genes and genomes as our reference gene lists.

#### Deconvolution of BULK RNA-Seq

Cibersortx is a tool which allows "in silico flow cytometry" and imputation of gene expression profiles from single cell data to provide an estimation of the abundances of member cell types in a mixed cell population of bulk RNA-Seq data.(55) Using the “cell fractions” module to allow deconvolution of bulk RNA-Seq data, we first generated a reference file using signature genes derived from the single cell transcriptomes, with the quantile normalisation option disabled. Average gene expression threshold (in log2 space) for cells with the same cellular phenotype was set to 0. The “replicates” option was set to 5 and “sampling” to 0.5. During the “Impute Cell Fractions” phase, S-mode batch correction was used, with quantile normalization again disabled and 500 permutations used for significance analysis.

This was then used to infer the cell fractions in the bulk mixture over 500 using the S-mode batch correction.

#### Single cell classification validation in publicly available datasets

To further verify our senescent cell classification in publicly available data (in supplementary figure 5), we used scPRED(30) a supervised classification method based on singular value decomposition and a support vector machine model. Using our R-UUO dataset as a reference, we trained our classifier using a Mixture Discriminant Analysis model and the default parameters, which returned high sensitivities and specificities including values >.9 for senescent epithelial cells. This was then used to classify cells in other datasets. DEG and pathway analysis was then performed as previously described.
